# Aging-related inflammation driven by cellular senescence enhances NAD consumption via activation of CD38^+^ pro-inflammatory macrophages

**DOI:** 10.1101/609438

**Authors:** Anthony J. Covarrubias, Abhijit Kale, Rosalba Perrone, Jose Alberto Lopez-Dominguez, Angela Oliveira Pisco, Herbert G. Kasler, Mark S. Schmidt, Christopher D. Wiley, Shankar S. Iyer, Natan Basisty, Qiuxia Wu, Ryan Kwok, Indra Heckenbach, Kyong-Oh Shin, Yong-Moon Lee, Issam Ben-Sahra, Melanie Ott, Birgit Schilling, Katsuhiko Ishihara, Stephen R. Quake, John Newman, Charles Brenner, Judith Campisi, Eric Verdin

## Abstract

Decline in tissue NAD levels during aging is linked to aging and its associated diseases. However, the mechanism for aging-associated NAD decline remains unclear. Here we report that pro-inflammatory M1-like macrophages, but not naïve or M2 macrophages, accumulate in metabolic tissues including visceral white adipose tissue and the liver during aging. Remarkably, these M1-like macrophages highly express the NAD consuming enzyme CD38 and have enhanced CD38-dependent NADase activity. We also find that senescent cells progressively accumulate in visceral white adipose tissue during aging and that inflammatory cytokines found in the supernatant from senescent cells (Senescence associated secretory proteins, SASP) induce macrophages to proliferate and to express CD38. These results highlight a new causal link between visceral tissue senescence and tissue NAD decline during aging and represent a novel therapeutic opportunity targeting maintenance of NAD levels during aging.

## Introduction

Nicotinamide adenine dinucleotide (NAD) is a redox coenzyme central to energy metabolism and an essential cofactor for non-redox NAD-dependent enzymes including sirtuins and poly-ADP-ribose polymerases (PARPs) ^1^. Early in the 20^th^ century, NAD deficiency was linked to pellagra, a disease characterized by dermatitis, diarrhea, dementia, and eventually death ^2^. More recently, a progressive decrease in NAD levels during aging in both rodents and humans has been documented in multiple tissues including the liver, adipose tissue, and muscle ^3^. Remarkably, restoration of NAD levels with the NAD precursor vitamins nicotinamide riboside (NR), nicotinamide (NAM) and nicotinic acid (NA), in addition to the biosynthetic NAD precursor nicotinamide mononucleotide (NMN) appear to mitigate aging-associated diseases ^3–5^. These observations have stimulated a flurry of research activity aiming to better understand how NAD levels affect the aging process and how or why NAD levels decline during aging, with the goal of developing therapeutics to combat aging-related disease.

NAD can be synthesized from tryptophan through the *de novo* pathway and by salvage of one of the three NAD precursor vitamins ^5^. While dietary precursors can contribute to NAD pools in a manner that depends on which pathways are expressed in particular tissues ^6^, the prevailing thought is that the recycling of NAM via nicotinamide phosphoribosyltransferase (NAMPT) is the predominant metabolic pathway most cells use to maintain intracellular NAD levels ^7^. The rate of NAD synthesis is countered by the rate of consumption by NAD consuming enzymes including sirtuins, PARPs, and the CD38 and CD157 NAD hydrolases. Importantly, the relative contribution of reduced de novo NAD biosynthesis, reduced NAM salvage, enhanced NAD consumption, or a combination of these effects to the NAD decrease observed during aging has not been established.

There is evidence of reduced NAM salvage during aging as aging-associated inflammation leads to reduced expression of NAMPT ^8^. There is also evidence for increased expression of key NAD consuming enzymes such as PARPs and CD38 during aging^9^. PARPs, particularly the ubiquitously expressed PARP1 and PARP2, are activated as a result of DNA damage during aging ^10^. Clinical conditions associated with defective DNA repair such as Cockayne syndrome and Werner’s disease show increased PARP activation, NAD depletion and progeroid (premature aging) diseases ^11–13^.

A recent report has also highlighted increased expression of CD38 during aging in visceral white adipose tissue^14^. CD38 is a transmembrane protein that consumes NAD to form cyclic ADP-ribose (cADPR), ADP-ribose (ADPR) and NAM ^14^. Importantly, mice lacking CD38 (*Cd38* KO) were protected from age-related NAD decline and had enhanced metabolic health and SIRT3 dependent mitochondrial function, supporting the model that CD38 is the primary NAD-consuming enzyme in age-related NAD decline in this tissue ^14^. However, these data did not address which cells express CD38 in aged tissues and what mechanism(s) drive aberrant CD38 expression during aging.

CD38 is ubiquitously expressed by immune cells and was first identified as a T cell activation marker ^15,16^. Its expression increases during inflammatory conditions including colitis, sepsis, and HIV infection ^17–19^. Chronic low grade inflammation during aging, termed “inflammaging” ^20^, is a leading mechanism of aging-associated diseases and has been described as a significant risk factor for morbidity and mortality ^21^. Sustained activation of the immune system is energetically costly and requires sufficient amounts of metabolites to fuel effector immune functions ^22^. Thus, the immune system and metabolism are highly integrated. Despite this knowledge, it is unclear how age-related inflammation affects NAD metabolism and the aging process.

To study possible links between age-associated NAD decline and inflammation, we have focused on metabolic tissues including visceral adipose tissue and the liver. By studying these tissues during the switch from lean to obese conditions much has been learned about how inflammation influences metabolic homeostasis, and how nutrient status in turn influences immunity ^23^. As obesity develops, peripheral monocytes are recruited to adipose tissue by adipokines, leading to a progressive infiltration of pro-inflammatory M1-like macrophages, and the displacement of anti-inflammatory resident M2 macrophages ^24–26^. Increased M1-like macrophages promote a pro-inflammatory state accompanied by insulin resistance, reduced rates of lipolysis, reduced tissue homeostasis, and the recruitment of macrophages to dying or stressed adipocytes resulting in the formation of “crown like structures” ^27,28^. This model suggests that macrophages, which are found in high abundance in key metabolic tissues such as liver and adipose tissue, may regulate metabolic health and disease through M1/M2 like-polarization and inflammatory signaling. We hypothesized that expression of NAD consuming enzymes including CD38 by resident-tissue macrophages contributes to their role in aging-associated metabolic changes by regulating NAD homeostasis.

Here, we report that pro-inflammatory M1 macrophages show increased CD38 expression, enhanced NADase activity, and production of NAD degradation byproducts NAM and ADPR. Using macrophages derived from WT and *Cd38* KO mice, we show that the high NADase activity of M1 macrophages is completely dependent on CD38 and not on other NAD consuming enzymes. We report that CD38 expression is elevated in resident macrophages from epididymal white adipose tissue (eWAT) and in the liver of old mice compared to young mice. Lastly, we show that enhanced CD38 expression by tissue resident macrophages during aging is driven by the senescence associated secretory phenotype (SASP), the secretion of inflammatory factors by senescent cells^29^. As senescent cells progressively accumulate in adipose tissue and liver during aging, these results highlight a new causal link between visceral tissue senescence and tissue NAD decline during aging. Thus, suppressing CD38 expression and NADase activity by targeting macrophages and senescent cells may provide a therapeutic target to help restore NAD levels during aging.

## Results

### M1 macrophage polarization leads to increased expression of the NAD hydrolases CD38 and CD157, and is characterized by enhanced degradation of NAD

Despite the recent emergence and renewed interest in both NAD metabolism and immuno-metabolism, little is known how NAD levels are regulated by immune cells and whether NAD levels influence immune cell function. Therefore, to better understand how tissue resident macrophages contribute to their role in aging-associated NAD metabolic changes, we first decided to survey the gene expression of enzymes that consume or are involved in the biosynthesis of NAD during pro and anti-inflammatory macrophage polarization. To test if different subsets of polarized macrophages have differential expression of NADase enzymes, we polarized primary mouse bone marrow-derived macrophages (BMDMs) to the classical pro-inflammatory M1 macrophage with the bacterial endotoxin LPS and to the anti-inflammatory alternative activated M2 macrophage with the Th2 cytokine IL-4. We observed that the only NADase enzymes significantly upregulated were *Cd38* (600-fold increase) and to a lesser degree its homologue *Cd157(Bst1)* (10-fold increase), while *Sirt1, Parp1* and *Parp2* were not significantly upregulated in M1 macrophages relative to naïve M0 and M2 macrophages (Figure 1A). A similar increase in *CD38* expression was also observed in human M1 macrophages (Figure S1A). This data suggest pro-inflammatory M1 macrophages may exhibit higher NADase activity, compared to M2 or untreated M0 macrophages. To test this hypothesis we utilized Nicotinamide 1,N^6^-etheno-adenine dinucleotide (ε-NAD), a modified version of NAD in which cleavage by an NAD hydrolase, such as CD38 or CD157, yields an increased fluorescence at 460nm that can be monitored over time. Incubating cellular lysates from untreated naïve (M0) or M2 macrophages showed no significant increase in fluorescence (Figure 1B). In contrast, M1 macrophage cellular lysates showed significantly increased fluorescence, consistent with increased NADase activity (Figure 1B and 1C).

**Figure 1.**
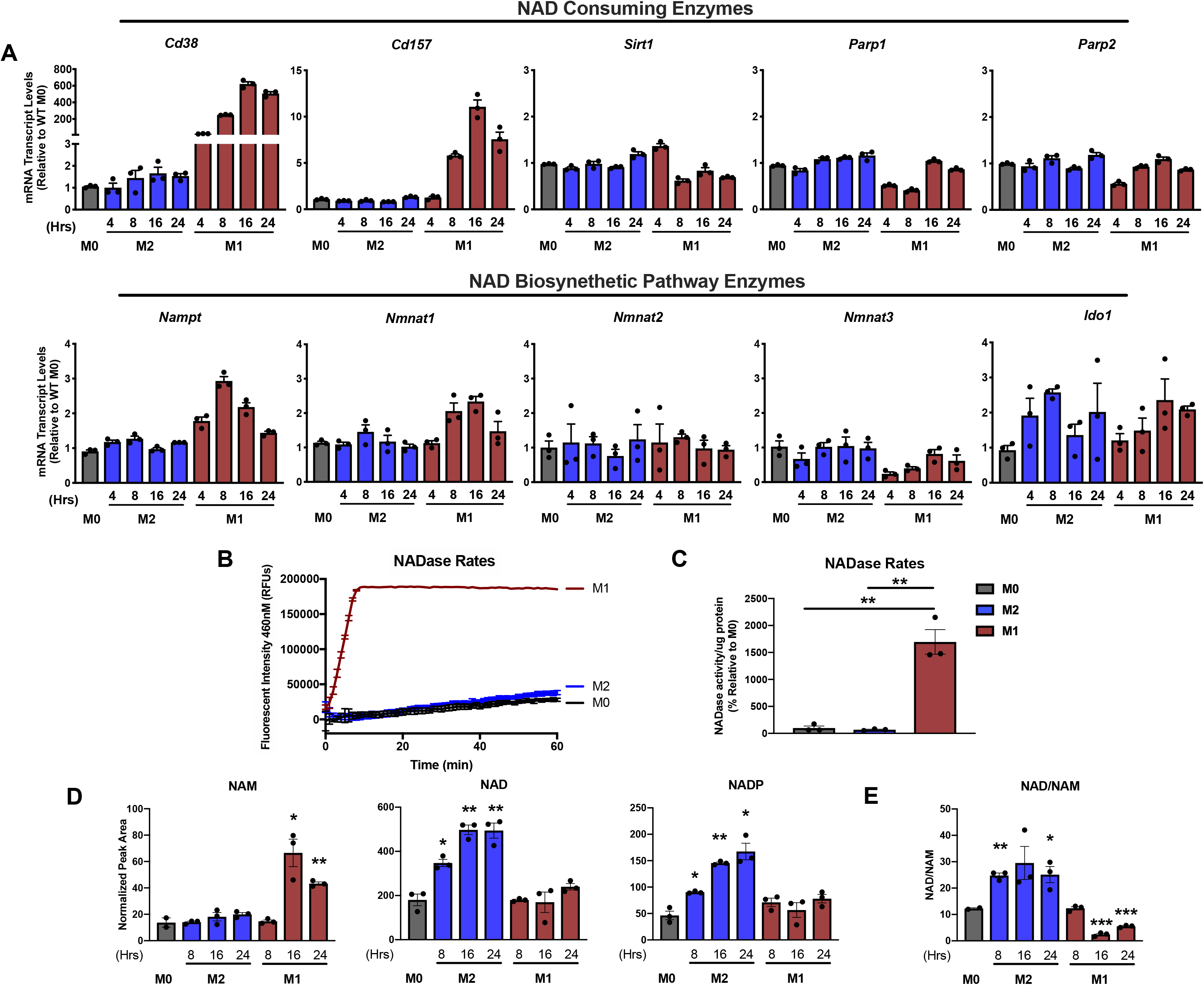
M1 macrophages are characterized by increased NADase activity. (A) mRNA expression of NAD consuming enzymes and NAD biosynthetic enzymes were measured using qPCR in untreated BMDMs (M0) or BMDMS treated with IL-4 (M2) or LPS (M1) for the indicated times. (B) NADase rates were measured in cell lysates from naive BMDMs (M0) or BMDMs treated with IL-4 (M2) or LPS (M1) after 16 hours of activation. (C) Quantification of the NADase activity rate. (D) LC-MS was used to quantify NAD and NAD related metabolites in naïve M0, M1, and M2, BMDMs for the indicated times. (E) NAD/NAM ratios from LC-MS data above for each indicated time point Data is showing the mean ± SEM. (n×3 biologically independent experiments). Statistical significance defined as \**P*<0.05, \*\**P*<0.01, and \*\*\**P*<0.00; two-sided Student’s t-test.

Phenotypes in mouse and human macrophages can differ significantly ^30^. To test whether these observations applied to human macrophages, we measured NADase activity in M0, M1 and M2 macrophages derived from peripheral blood monocytes. We observed a similar increase in NAD hydrolysis in human M1 macrophages when compared to naïve and M2 subsets (Figure S1B). Taken together our data shows that enhanced NAD hydrolysis is a highly conserved feature of inflammatory M1 macrophages in both human and mice.

NAD is a critical regulator of metabolic pathways since it is the major hydride acceptor and co-factor for metabolic enzymes in glycolysis, TCA cycle, and glutaminolysis. Nutrient sensing and metabolic pathways are critical for macrophage activation, gene transcription, and other effector functions ^31,32^. The change in NAD hydrolase activity shown above suggested that different macrophage subtypes might exhibit different NAD levels. No reports have investigated how NAD levels might be linked to the metabolic reprogramming of macrophage metabolism or how NAD levels are affected in different macrophage polarization states. To address this question, we used targeted LC-MS to quantify the NAD metabolome in polarized macrophages. Importantly, other NAD-consuming enzymes such as PARPs, sirtuins, CD38 and CD157 all produce NAM as a byproduct of NAD hydrolysis. Consistent with the high NADase activity of M1 macrophages, we observed significantly elevated levels of NAM in M1 macrophages, but not in naïve M0 and M2 macrophages (Figure 1D). We also observed elevated levels of NAD and NADP in M2 macrophages but not in M1 macrophages (Figure 1D). Furthermore, NAD/NAM ratios were significantly lower in M1 macrophages (Figure 1E).

Despite the high NADase activity associated with M1 macrophages, their NAD levels seemed to be stably maintained compared to untreated naive macrophages (Figure 1D). This suggested that M1 macrophages might upregulate their biosynthetic and NAM salvage pathways to compensate for increased NAD degradation. The tissue culture medium used in our experiments lacks nicotinic acid but contains NAM. Under these conditions, NAD can be derived *de novo* from tryptophan or recycled from NAM via the NAM salvage pathway. Therefore, we measured mRNA levels of indoleamine 2,3-dioxygenase 1 (*Ido1)*, along with other key enzymes in the *de novo* NAD synthetic pathway (Figure 1A, S1C-D), and the NAM salvage pathway enzymes *Nampt* and nicotinamide mononucleotide adenylyltransferases 1-3 (*Nmnat1-3*) (Figure 1A). In M1 macrophages, we observed an increase in mRNA levels for *Nampt* and *Nmnat1*, which maintain nuclear NAD levels (Figure 1A). We did not see any significant changes in *Nmnat2* or *Nmnat3*, which regulate NAM salvage in the cytoplasm or mitochondria respectively, or in *Ido1* between M1 and M2 macrophages (Figure 1A). Interestingly, the gene expression of several key enzymes in the de novo synthesis pathway were downregulated during M1 polarization, including kynureninase (*Kynu*) and 3-hydroxy-anthranilic acid oxygenase (*Haao*) (Figure S1D). Furthermore, using LC-MS we observe increased levels of tryptophan, kynurenine and kynurenic acid in M1 macrophages compared to M0 and M2 macrophages (Figure S1E). However, we were unable to detect other tryptophan pathway metabolites downstream of kynurenine in either M0, M1 or M2 macrophages, which suggests, but does not prove, that tryptophan may not significantly contribute to *de novo* NAD synthesis in macrophages regardless of their differentiation stage.

### The NAM salvage pathway is a critical regulator of NAD levels during M1 and M2 macrophage polarization and fine tunes macrophage gene expression

Our observation that *de novo* synthesis of NAD from tryptophan may not significantly contribute to maintenance of NAD levels in M1 and M2 macrophages predict a critical role for the NAM salvage pathway in maintenance of NAD levels in macrophages. Indeed, we observed that macrophages treated with FK866, a specific inhibitor of NAMPT (Figure 2A) showed a near complete suppression of NAD levels both in M1 and M2 macrophages (Figure 2B). Our data indicate that macrophages rely almost exclusively on the NAM salvage pathway to sustain their NAD levels and that M1 macrophages compensate for their enhanced NADase activity by increasing mRNA expression of key NAD salvage pathway enzymes, *Nampt* and *Nmnat1* (Figure 1A).

**Figure 2.**
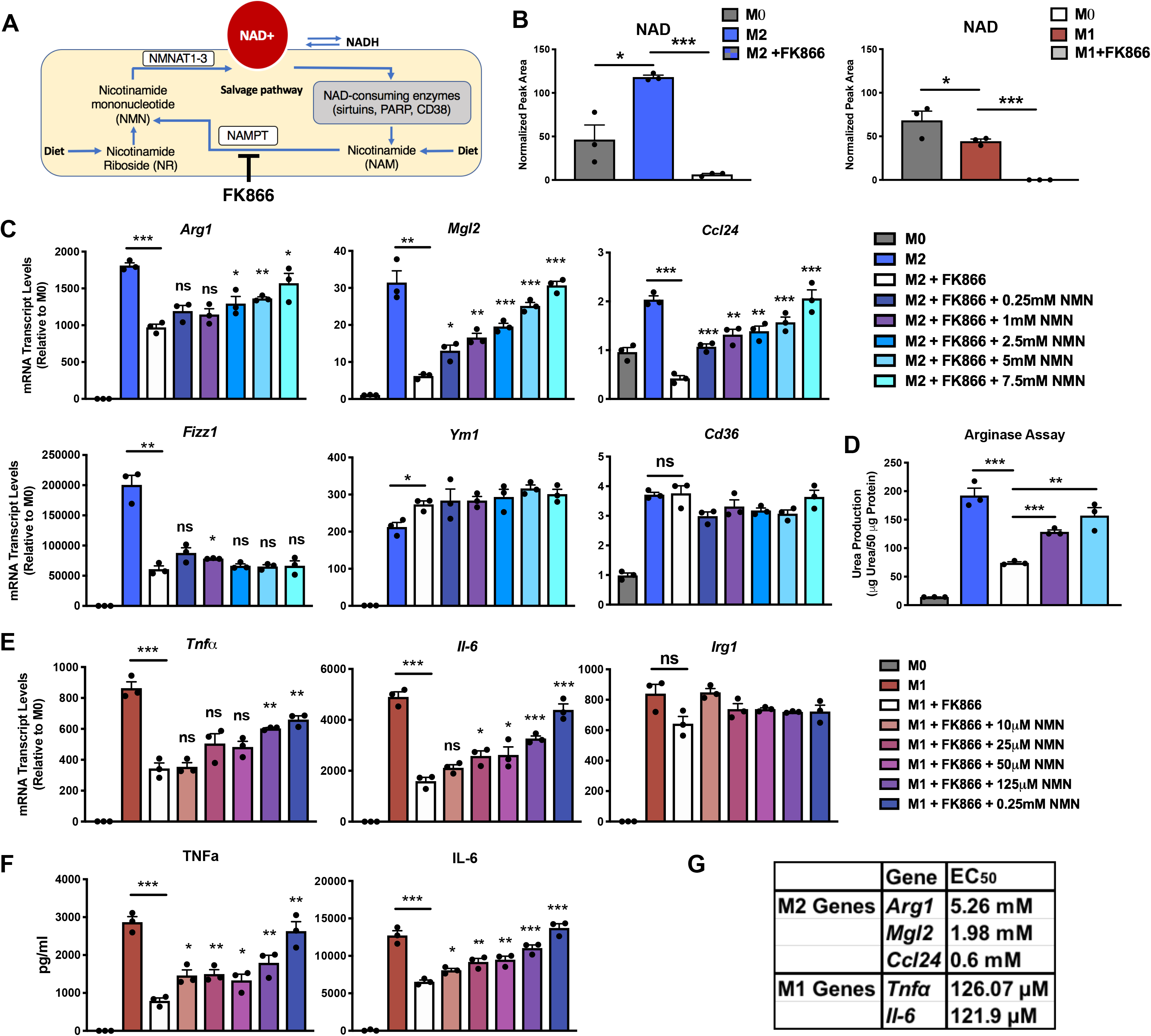
The NAM salvage pathway controls NAD levels to regulate macrophage activation and polarization. (A) Schematic representation of the NAM salvage pathway. (B) NAD levels were measured by LC-MS in M0, M1 and M2 BMDMs pre-treated with or without 10nM/ml FK866 for 6 hours prior to stimulation with LPS for 6 hours or IL-4 for 16 hours. (C) mRNA levels of M2 genes in BMDMs pre-treated with or without FK866 and NMN for 6 hours prior to stimulation with IL-4 for 16 hours. (D) Arginase assay of M2 macrophages pre-treated with or without FK866 and NMN for 6 hours prior to stimulation with IL-4 for 16 hours. (E-F) mRNA levels and ELISA measurements of TNF-α and IL-6 in supernatants from BMDMs pre-treated with or without FK866 and NMN for 6 hours prior to stimulation with LPS for 6 hours. (G) Quantification of the half maximal effective concentration (EC_50_) for NMN used to rescue M1 and M2 macrophage gene expression. Data is showing the mean ± SEM. (n=3 biologically independent experiments). Statistical significance defined as \**P*<0.05, \*\**P*<0.01, and \*\*\**P*<0.00; two-sided Student’s t-test. All statistical comparisons are relative to M2/M1 + FK866.

In the experiments described above, we consistently observed rising NAD levels during M2 polarization compared to M1 and M0 macrophages (Figure 1D and 2B). This result suggested that M2 polarization requires higher NAD levels. To test if NAD levels can regulate macrophage polarization, we pre-treated BMDMs with the NAMPT inhibitor FK866, to lower NAD levels, prior to treatment with IL-4. This led to decreased mRNA expression of multiple canonical M2 markers (*Arg1*, *Mgl2*, *Ccl24* and *Fizz1*) (Figure 2C), however this was not universal since the M2 markers *Ym1* and *Cd36* were unaffected. In agreement with reduced expression of the gene *Arg1*, arginase activity, a key anti-inflammatory effector function of M2 macrophages, was also attenuated by FK866 (Figure 2D). We have previously shown that the same subset of FK866 sensitive M2 genes (*Arg1*, *Mgl2*, *Ccl24*, and *Fizz1*) are fine-tuned and integrated to metabolic inputs, while *Ym1* and *Cd36* are insensitive to metabolite levels including amino acids ^33^. To confirm the role of NAD depletion, we supplemented macrophages treated with FK866 with different doses of nicotinamide mononucleotide (NMN) or nicotinamide riboside (NR) in the culture medium. Both NMN and NR are NAD precursors that feed into the NAM salvage pathway downstream of NAMPT (Figure 2A). Both rescued the mRNA expression of M2 genes inhibited by FK866 (Figure 2C and S2A) and arginase activity (Figure 2D) in a dose-dependent manner, with the exception of *Fizz1*. In contrast to a previous report ^34^, we observed that pre-treatment of macrophages with FK866 also suppressed mRNA expression and secretion of M1 inflammatory cytokines such as *Il-6* and *Tnfα* but not the metabolic enzyme *Irg1* (Figure 2E and 2F). NMN and NR also restored mRNA expression and protein levels of inflammatory cytokines in M1 macrophages (Figure 2E, 2F, and S2B). However, much higher concentrations of NMN (Figure 2C-F) and NR (Figure S2A and S2B) were needed to rescue FK866-sensitive M2 genes compared to M1 genes (Figure 2G).

These results show that the NAM salvage pathway regulates NAD levels in macrophages, and that NAD is a critical regulator of macrophage polarization and regulates the mRNA abundance of a subset of M2 and M1 genes. Importantly, we show that M1 and M2 macrophages have different requirements for NAD levels, with M2 macrophages requiring higher levels of NAD to support M2 gene expression and immune effector functions (Figure 2G). Thus, we provide evidence that the NAM salvage pathway is a critical regulator of macrophage activation and polarization in both the M1 and M2 state, and fine tunes the gene expression of a subset of M1 and M2 genes.

### The NAD consuming enzyme CD38 is exclusively expressed by M1 macrophages and is the primary enzyme responsible for enhanced NAD hydrolase activity observed in M1 macrophages

In Figure 1, we observed that M1 macrophage polarization is characterized by enhanced NADase activity and correlated with enhanced expression of the NAD hydrolases *Cd38* and its homologue *Cd157* (*Bst1*). To test the hypothesis that the high NADase activity observed in M1 macrophages is CD38/CD157-dependent, we first analyzed CD38 surface expression using flow cytometry in BMDMs. We compared M0, M1 and M2 macrophages obtained from wild type mice (WT) vs mice lacking CD38 (*Cd38* KO) ^35^. Consistent with the high *Cd38* mRNA levels in M1 macrophages (Figure 1A), we found CD38 to be exclusively expressed by WT M1 macrophages (Figure 3A, 3B and S3A), a finding also reported in a recent publication^36^. In contrast, PARP and sirtuin expression is unaffected in *Cd38* KO macrophages or during M1 and M2 polarization (Figure 3B). In support of our hypothesis that NADase activity observed in M1 macrophages is CD38-dependent, we found cell lysates from M1 macrophages lacking CD38 showed near basal NADase activity compared to WT M1 macrophages (Figures 3C and 3D). These data indicate that NADase activity of M1 macrophages specifically requires CD38, and not CD157 or any other NAD-consuming enzyme. CD38 can be localized to cell membranes in a type II and type III orientation, with its enzymatic domain facing outside (type II) or inside the cell (type III) ^37–39^. To test whether the CD38 enzymatic activity was predominantly localized to the extracellular vs. intracellular space, we measured NAD degradation of the non-cell permeable εNAD in intact cells. We found that extracellular εNAD hydrolysis occurred only in M1 macrophages (Figure S3B), although to a lesser extent than what we observed in cell lysates (Figures 3D). These data demonstrate that CD38 cleaves NAD outside and inside cells and that a significant fraction of CD38 NADase activity in macrophages is intracellular.

**Figure 3.**
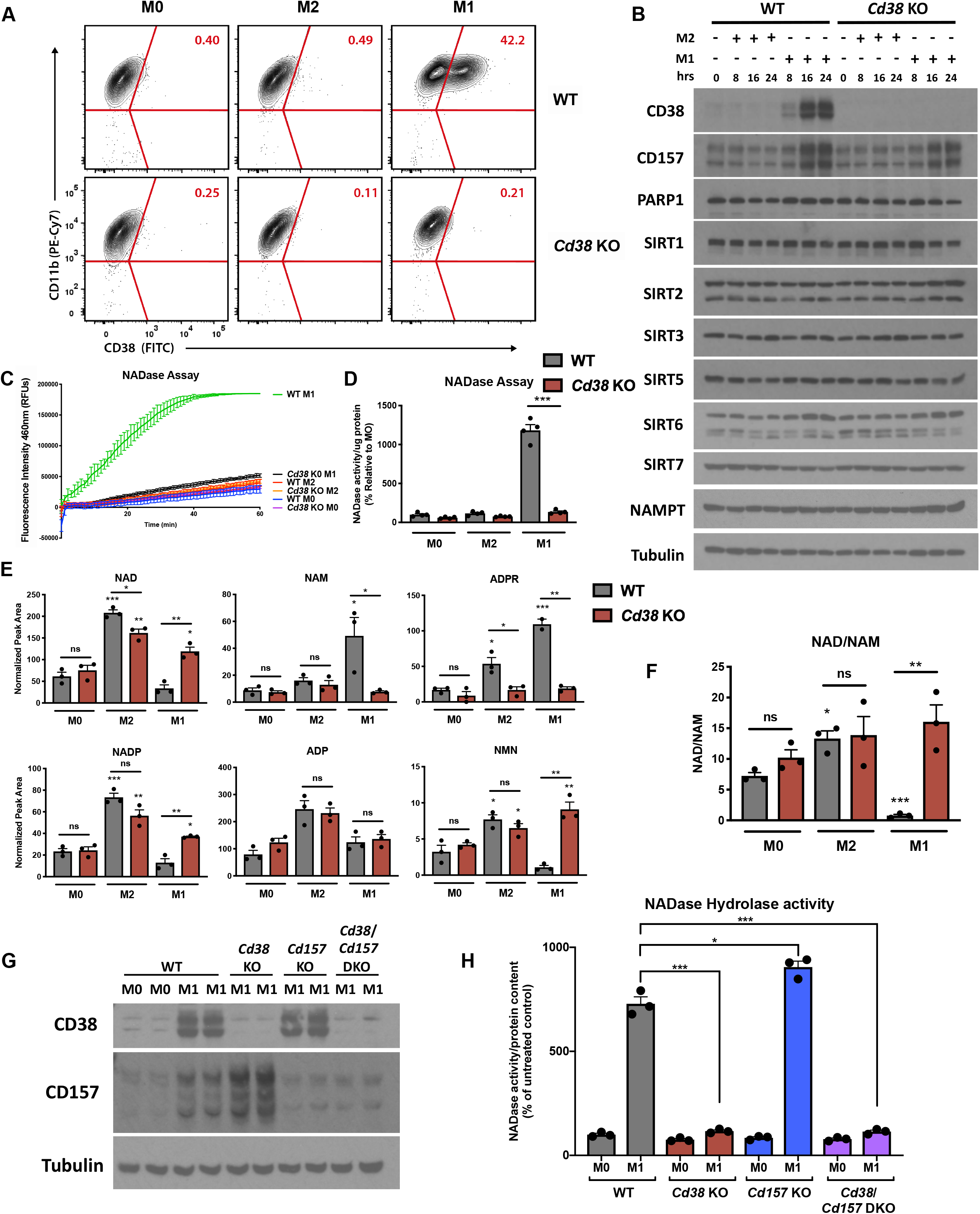
The high NADase activity in M1 macrophages is CD38 dependent. (A) Representative flow cytometry plots comparing CD38 surface staining in naive (M0) WT and *Cd38 KO* BMDMs or treated with IL-4 (M2) and LPS (M1) for 16 hours. (B) Western blot of NADase enzymes in M0, M1, and M2 WT and *Cd38 KO* for the indicated times (C) NADase activity rates measured in WT and *Cd38 KO* MO, M2, and M1 BMDMs activated for 16 hours. (D) Quantification of the NADase activity rate. (E) LC-MS was used to quantify NAD and NAD related metabolites in M0, M2, and M1 WT and *Cd38 KO* BMDMs activated for 16 hours. (F) NAD/NAM ratios from LC-MS data above. (G)Western blot of CD38 and CD157 enzymes in MO and M1 BMDMs from WT, *Cd38* KO*, Cd157* KO*, and Cd38/Cd157* DKO mice stimulated 16 hours. (H)NADase activity rates measured in in MO and M1 BMDMs from WT, *Cd38* KO*, Cd157* KO*, and Cd38/Cd157* DKO mice stimulated 16 hours. Data is showing the mean ± SEM. (n=3 biologically independent experiments, n=4 in D). Statistical significance defined as \**P*<0.05, \*\**P*<0.01, and \*\*\**P*<0.00; two-sided Student’s t-test. Unless noted with a bar, all statistical comparisons are relative to untreated WT or *Cd38* KO sample.

To further investigate how CD38 regulates intracellular NAD levels in macrophages, we quantified NAD and its metabolites in WT and *Cd38* KO BMDMs. We found that WT M1 macrophages had increased levels of NAM and ADP Ribose (ADPR), two specific products resulting from CD38 activity on NAD (Figure 3E). Importantly, the production of both metabolites was suppressed to basal levels in M1 macrophages derived from *Cd38* KOs (Figure 3E). *Cd38* KO M1 macrophages also had significantly higher levels of NAD and NADP and a higher NAD/NAM ratio compared to WT M1 macrophages (Figure 3E and 3F). Additionally, we found no significant difference in the level of the nucleotide ADP in WT vs *Cd38* KO M1 macrophages (Figure 3E). Therefore, CD38 specifically consumes and regulates NAD metabolite levels and does not affect the levels of other adenine-containing nucleotides. These data support the model that CD38 is a key regulator of NAD and its metabolites NAM and ADPR in M1 macrophages. These results seem to exclude a major role for CD157, whose mRNA and protein levels also increased in M1 macrophages (Figure 3B and 3G), as an NAD hydrolase. Indeed, while *Cd38* KO and *Cd38/Cd157* Double KO (DKO) M1 macrophages have attenuated NADase activity, *Cd157* KO M1 macrophage still have elevated NADase activity similar to WT M1 macrophages (Figure 3G-H). Interestingly, recent reports suggest the preferred CD157 substrate is the NAD precursor NR ^40,41^. In indirect support of this, we found that CD157 expression which was enhanced in response to LPS treatment was associated with significantly reduced NR levels in both WT and *Cd38* KO M1 macrophages (Figure S3C and S3D). Thus, although CD157 may indirectly influence NAD levels in M1 macrophages by consuming NR, the enhanced NADase activity of pro-inflammatory M1 macrophages is mediated by CD38.

### Aging is associated with enhanced adipose tissue inflammation and the accumulation of CD38 positive macrophages

To verify that NAD levels decline in adipose tissue during aging, we measured NAD levels in epididymal white adipose tissue (eWAT) of mice at 6 months (young) and 25 months (old). We observed close to 50% reduction of NAD and NADP in old mice compared to young (Figure 4A), consistent with previous reports ^8,14^. Our results above showing that M1 macrophages are characterized by enhanced CD38 dependent NAD degradation suggest that M1 macrophages may be a key driver of NAD loss during conditions of chronic-inflammation, such that occurs during aging. To test whether declining NAD levels during aging are associated with chronic-inflammation, we measured gene mRNA expression analysis of old vs young eWAT. Our analysis revealed increased expression of *Cd38*, senescence markers (*p16*^*Ink4a*^, *p21*^*Cip1*^), and SASP genes including inflammatory cytokines and chemokines (*Il-6, Il-1β, ll-1α, Il-10* and *Cxcl1*) in aged eWAT when compared to that in young mice (Figure 4B and S4A). Conversely, M2 macrophage markers trended down (such as *Arg1*, *Fizz1* and *Mgl2*) or were unchanged (*Ccl24*) (Figure 4B). Furthermore, our analysis of eWAT from old and young mice revealed increased mRNA transcript and protein expression of the macrophage marker CD68 in old mice, consistent with greater macrophage content in old tissues (Figure 4B and 4C). Interestingly, old eWAT tissue were also characterized by greater protein expression of the NAD hydrolases CD38 and CD157 (Figure 4C and 4D), and significantly elevated levels of inflammasome activation as indicated by higher levels of the 20 kDa processed form of Caspase-1 (Figure 4C), which cleaves and activates the inflammatory cytokines IL-1β and IL-18.

**Figure 4.**
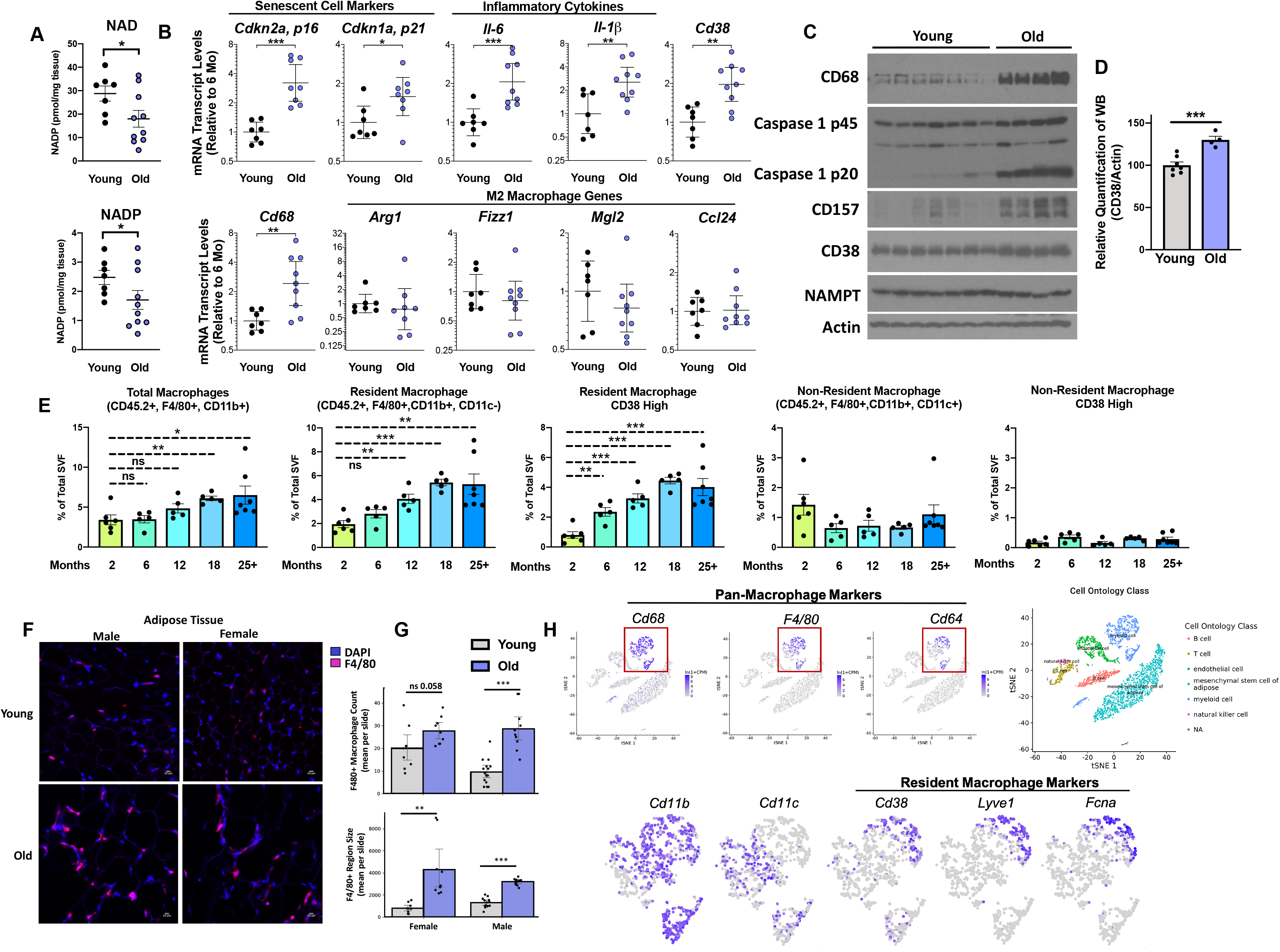
NAD decline during aging is associated with increased CD38+ tissue-resident macrophages in eWAT. (A) LC-MS was used to quantify NAD and NADP in whole visceral epididymal white adipose tissue (eWAT) from 6-month and 25-month-old mice. NAD and NADP concentrations are shown as pmol/mg of tissue. (B) RNA levels of senescence markers, inflammatory genes, macrophage marker *Cd68*, and M2 genes in whole eWAT from young (6-month) and old (25-month) mice. (C) Western blot of the indicated target proteins in whole eWAT from young (3-month) and old (30-month) mice. (D) ImageJ quantification of CD38 protein expression relative to Actin in whole eWAT from young (3-month) and old (30-month) mice. (E) Quantification of total macrophages, CD38+ resident macrophages, and CD38+ non-resident macrophages isolated from eWAT of mice for the indicated age. (F) Immunofluorescence (IF) of the macrophage marker/antigen F4/80 (magenta) and nuclei with DAPI (Blue) in eWAT in young (4-month) and old (26-month) male and female WT mice. Scale bars represents 10μm. Representative of 4-5 mice/group. (G) Trained neural network analysis of IF images quantifying the mean number of macrophages (defined as F4/80 colocalized to DAPI) per slide, and mean F4/80+ region size for old and young eWAT graphed as mean per slide. (H) Analysis of CD38 and other macrophage markers in eWAT from single-cell transcriptome data using the Tabula-Muris database (https://tabula-muris.ds.czbiohub.org). Data from individual mice are shown for *in vivo* experiments. Data is showing the mean ± SEM. Statistical significance defined as \**P*<0.05, \*\**P*<0.01, and \*\*\**P*<0.00; two-sided Student’s t-test except for 4A one-sided t-test was used.

Next, we analyzed CD38 expression in macrophages, the predominant immune cell in visceral adipose tissue ^42^, isolated from the stromal vascular fraction (SVF) of eWAT from old and young mice. As previously reported, flow cytometry analysis of the SVF can be used to distinguish resident and non-resident populations of adipose tissue macrophages based on cell surface markers expressed by macrophages, including F4/80 (EMR1) and CD11b (ITGAM) which are expressed by both resident and non-resident adipose tissue macrophages, and the integrin CD11c (ITGAX) which is expressed by non-resident adipose tissue macrophages ^42,43^. Using a similar gating strategy as Cho et al.^43^, and outlined in Figure S4B, we found a significant increase in the proportion of macrophages in the SVF of adipose tissue particularly after 12 months of age (Figure 4E). Additionally, we used immunofluorescent (IF) staining of the macrophage specific marker F4/80, coupled with an unbiased computational analysis using a trained neural network to quantitate the amount of F4/80+ macrophages in both old and young adipose tissue. Visually and with a trained neural network we were able to detect greater abundance of F4/80+ macrophages in both old female and male mice compared to younger mice (Figure 4F and 4G). This increase in the number of total macrophages in aged tissue was accounted for by an increase in the proportion of macrophages bearing a signature of resident macrophages (F4/80+, CD11b+, CD11c-), but not in the non-resident macrophage population (F4/80+, CD11b+, CD11c+) (Figure 4E). Interestingly, resident macrophages from older animals, but not the non-resident macrophages, showed progressively increased CD38 expression (Figure 4E and S4C).

To further investigate in an unbiased fashion what macrophage populations express CD38 in adipose tissue we used the Tabula Muris single-cell RNA-Seq database^44^, a powerful tool to dissect gene expression in single cell populations in multiple organs and tissues. We found that CD38 expression was specific to adipose tissue resident macrophages (CD68+, CD64+, F4/80+, CD11b+, CD11c−) and not the non-resident population (CD68+, CD64+, F4/80+, CD11b+, CD11c+) (Figure 4H), consistent with our flow cytometry analysis (Figure 4E).

Furthermore, recent manuscripts have characterized the gene expression of tissue-resident immune cell populations and found that the genes *Lyve1* and *Fcna* are co-expressed along with CD38 by a subset of tissue-resident macrophages^45,46^, this finding was also confirmed in adipose tissue macrophages using the Tabula Muris database (Figure 4H). Our previous data (Figure 1 and 3) suggest CD38 is a marker of inflammatory macrophages. Thus, our data suggest that during aging, resident adipose tissue macrophages are polarizing from a M2 to M1-like phenotype. A similar switch from M2 to M1 macrophages has been reported during the metabolic stress of obesity ^24^. Although CD38 is also expressed by B cells and activated T cells, we did not see a significant increase in CD38 expression during aging in non-macrophage immune cells in the SVF (Figure S4C). Thus, compared to young mice, we found adipose tissue from old animals to be significantly more inflamed, have increased accumulation of senescent cells and macrophages, and a noticeable shift in tissue-resident macrophage polarization from a M2 to a pro-inflammatory M1-like state characterized by increased CD38 expression.

### Microbial products and the senescence associated secretory phenotype promotes macrophage CD38 expression

Increased CD38 expression and enhanced NADase activity occurs in response to LPS-driven activation of toll-like receptor 4 (TLR4) in macrophages (Figure 1, 2, and 3) as a programmed immune response to gram negative bacteria ^47^. A recent study showed that gut intestinal barrier permeability increases during aging leading to significantly higher levels of endotoxins, including LPS, in aging tissues ^48,49^. To test whether other TLR ligands besides LPS can increase *Cd38* mRNA expression in macrophages, we treated BMDMs with multiple pathogen associated molecular patterns (PAMPs) and danger associated molecular patterns (DAMPs) that act as TLR ligands (Figure 5A and 5B). All PAMP TLR ligands highly activated *Cd38* mRNA expression, with the exception of poly(I:C), a synthetic double stranded RNA that mimics viral RNA which significantly induced *Cd38* expression but more modestly (Figure 5A). To further explore whether exposure to low amounts of endotoxins can promote CD38 expression *in vivo* and affect NAD levels, we analyzed eWAT from mice treated for four-weeks intraperitoneally with a non-lethal dose of LPS or PBS as a control. Similar to aged mice (Figure 4B and 4C), in LPS treated mice we observed significantly elevated transcript levels of inflammatory cytokine expression (*Tnfα* and *Il-1β*), greater expression of the macrophage markers *Cd68* and *F4/80*, indicative of greater macrophage abundance, and higher expression of the NAD hydrolases C*d38* and *Cd157* (Figure 5C). In contrast the gene expression of other NAD consuming enzymes including sirtuins and PARPs were unchanged or significantly down-regulated in LPS treated mice (Figure S5A). Metabolomic analysis of the eWAT also revealed decreased NAD and NADP levels in the inflamed LPS treated eWAT compared to PBS treated mice (Figure 5D). Thus, our data provides evidence that exposure to low-levels of endotoxins and PAMPs is sufficient to promote a low-grade chronic inflammatory state associated with increased expression of CD38 and depletion of NAD levels.

**Figure 5.**
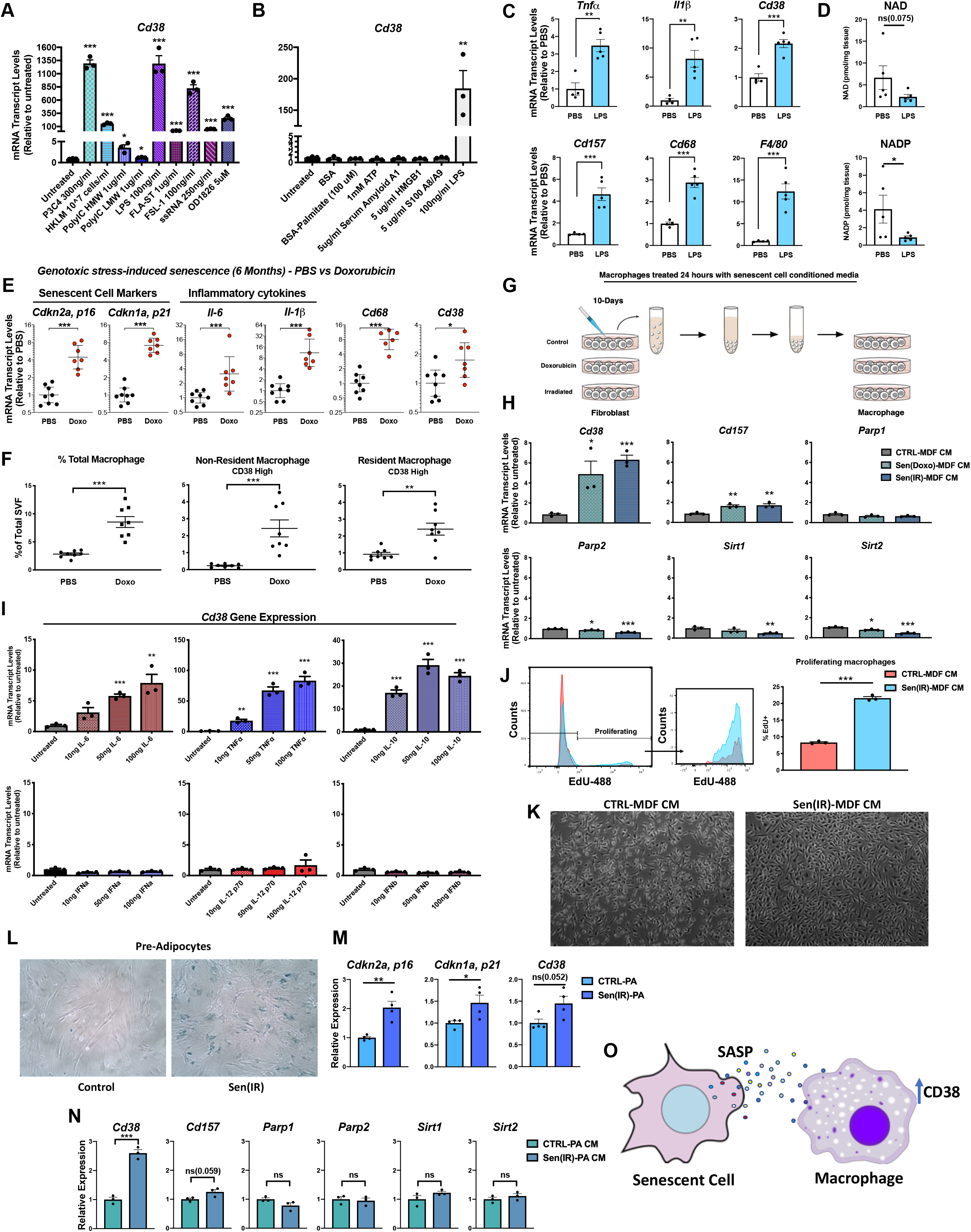
Secreted cytokines from senescent cells promotes CD38 expression and proliferation in macrophages. (A) mRNA levels of *Cd38* in BMDMs treated with the indicated TLR ligands for 16 hours. (B) mRNA levels of *Cd38* in BMDMs treated with the indicated DAMP ligands for 16 hours. (C) mRNA levels in eWAT from 3-month-old mice IP injected with PBS or LPS for 4-weeks. (D) LC-MS was used to quantify NAD and NADP in eWAT from 3-month-old mice IP injected with PBS or LPS for 4-weeks. NAD and NADP concentrations are shown as pmol/mg of tissue. (E) mRNA levels in eWAT from 6-month-old mice IP injected with PBS or doxorubicin. Doxorubicin increases the expression of senescence markers, inflammatory cytokines, macrophage marker *Cd68* and *Cd38* in total tissue and in isolated macrophages. (F) Flow cytometry analysis and quantification of CD38+ macrophages isolated from 6-month-old mice IP injected with PBS or doxorubicin. (G) Conditioned media (CM) was isolated from non-senescent control mouse dermal fibroblasts (CTRL-MDF), doxorubicin treated senescent MDFs (Sen(Doxo)-MDF), or irradiated senescent MDFs (Sen(IR)-MDF) 10-days post-treatment, and then used to stimulate BMDMs for 24 hours. (H) mRNA levels of *Cd38* and other NAD consuming enzymes in BMDMs treated for 24 hours with conditioned media (CM) from non-senescent control mouse dermal fibroblasts (CTRL-MDF), doxorubicin treated senescent MDFs (Sen(Doxo)-MDF), or irradiated senescent MDFs (Sen(IR)-MDF). (I) mRNA levels of *Cd38* in BMDMs treated with the indicated concentration (ng/ml) of recombinant mouse cytokines for 24 hours. (J) Flow cytometry of EdU+ BMDMs treated with Sen(IR)-mouse dermal fibroblast conditioned media (Sen(IR)-MDF CM) or non-senescent control mouse dermal fibroblast conditioned media (CTRL-MDF CM) for 24 hours. (K) Representative bright field microscopy image of BMDMs treated with CTRL-MDF CM or Sen(IR)-MDF CM for 24 hours. For *in vivo* experiments, data from individual mice are shown. (L) SA-β-Galactosidase staining in control (CTRL-PA) or irradiated senescent primary mouse pre-adipocytes (Sen(IR)-PA). (M) mRNA levels of the indicated genes in non-senescent control pre-adipocytes (CTRL-PA) or irradiated senescent primary mouse pre-adipocytes (Sen(IR)-PA). (N) mRNA levels of *Cd38* and other NAD consuming enzymes in BMDMs treated with conditioned media from non-senescent control preadipocytes (CTRL-PA CM) or irradiated senescent primary mouse pre-adipocytes (Sen(IR)-PA CM) for 24 hours. (O) Model showing that inflammatory cytokines (SASP) derived from senescent cells can promote macrophage expression of CD38. Data from individual mice are shown for *in vivo* experiments. Data is showing the mean ± SEM (n=3 biologically independent experiments, n=4 in M). Statistical significance defined as \**P*<0.05, \*\**P*<0.01, and \*\*\**P*<0.00; two-sided Student’s t-test except for 5D one-tailed t-test was used.

Senescent cells represent a key source of sterile inflammation in the aging process and are thought to play a key role in the chronic inflammation and chronic diseases associated with aging ^50^. Many stressors, including radiation, genotoxic chemotherapies, oncogene activation, replicative exhaustion and metabolic imbalances induce a senescence response. Senescent cells are characterized by cell cycle arrest, increased senescence-associated β-galactosidase activity, and by a complex senescence-associated secretory phenotype (SASP) that includes pro-inflammatory chemokines, DAMPs, and cytokines ^51^. Mounting evidence suggests that senescent cells can drive a surprisingly diverse array of aging phenotypes and diseases, largely through the SASP. In support of this model, eliminating senescent cells, using either transgenic mouse models or a new class of drugs termed senolytics, can help maintain homeostasis in aged or damaged tissues, and postpone or ameliorate many age-related pathologies, and even extend lifespan^52–54^.

Therefore, in addition to microbial endotoxins, we sought to investigate whether cellular senescence influences macrophage polarization and CD38 expression during aging. To test this hypothesis, we IP injected young mice with the chemotherapeutic agent doxorubicin. Doxorubicin intercalates into DNA and inhibits topoisomerase II activity, thereby inducing cellular senescence both in tissue culture and *in vivo* ^55,56^. In doxorubicin treated mice, mRNA expression of *Cd38* in eWAT was increased in parallel with senescence markers (*p16* and *p21*) and inflammatory cytokines and chemokines (Figure 5E and S5B), similar to that observed during aging (Figure 4B). Flow cytometry of macrophages extracted from eWAT from doxorubicin-treated mice showed increased macrophage accumulation compared to control mice (Figure 5F). Furthermore, doxorubicin treatment also led to increased CD38+ resident and non-resident macrophages (Figure 5F and S5C). These results support the model that senescent cells and the SASP represent a key mechanism for increased CD38 expression in adipose tissue macrophages found in aging eWAT.

To further test the hypothesis that SASP from senescent cells directly drives CD38 expression in macrophages, we induced senescence in mouse dermal fibroblasts by ionizing radiation (Sen(IR)-fibroblasts) or doxorubicin (Sen(Doxo)-fibroblasts) and purified conditioned medium from each of these conditions ^55,57^ (Figure 5G). Macrophages cultured for 24 hours in conditioned medium from senescent cells specifically upregulated *Cd38* and to a lesser extent *Cd157*, but not other NAD consuming enzymes including *Parp1*, *Parp2*, *Sirt1* and *Sirt2* (Figure 5H). Consistent with this observation, co-culturing WT macrophages with Sen(IR)-fibroblasts increased *Cd38* expression to a similar extent (Figure S5D). This did not occur in WT macrophages co-cultured with non-senescent fibroblasts or in *Cd38* KO macrophages (Figure S5D). Furthermore, we did not observe a significant increase in *Cd38* mRNA levels in Sen(IR)-fibroblasts alone compared to non-senescent fibroblasts (Figure S5D).

To investigate what components of the SASP can elicit CD38 expression in macrophages we first treated macrophages with DAMPs, endogenous alarmin molecules derived from damaged or dead cells, that have been reported to activate TLRs ^58^. Senescent human and mouse fibroblasts secrete nuclear protein HMGB1, a DAMP which has been reported to activate TLR4 in addition to RAGE (receptor for advanced glycation end products) to promote inflammatory signaling pathways ^59^. Surprisingly, treatment of BMDMs with multiple DAMPs including HMGB1 did not activate CD38 expression (Figure 5B), suggesting that individual DAMPs may not be a mechanism of CD38 activation in macrophages. In addition to DAMPs, cytokines such as TNFα and type 1 interferons (IFNα/IFNβ) have also been reported to drive CD38 expression in mouse and human macrophages ^60,61^. We observed that Sen(IR)-fibroblasts significantly up-regulate transcription of *Ifnα*, along with other inflammatory cytokines (*Tnfα, Il-12, Il-6,* and *Il-10*) (Figure S5E) compared to normal fibroblasts. Therefore, we treated BMDMs with these recombinant cytokines and measured *Cd38* mRNA transcript levels (Figure 5I). Interestingly, while IFNα, IFNβ, and IL-12 did not promote *Cd38* expression, TNF-α, IL-6, and IL-10 significantly increased *Cd38* expression (Figure 5I). Thus, despite being reported to activate *Cd38* expression, the inability of type-1 interferons to promote *Cd38* expression are consistent with poly(I:C) being a weak activator of *Cd38* expression compared to other TLR ligands (Figure 5E). The poly(I:C) receptor TLR3 is the only TLR whose signaling is MYD88-independent and signals through IRF3, which leads to upregulation of type 1 interferons and interferon inducible genes ^62^, but only weakly activates expression of *Cd38* (Figure 5A). Taken together these observations show that sterile sources of inflammation, including inflammatory cytokines found in the SASP are sufficient to promote the expression of *Cd38* in macrophages.

One of the most striking features we observed in aging eWAT is the accumulation of resident macrophages in mouse eWAT (Figure 4). Resident adipose tissue macrophages are seeded in tissues during development and have the capacity to self-renew, which is driven by CSF1 and other growth factors including MCP-1 and IL-4 during helminth infections ^63–67^. Interestingly, Sen(IR)-fibroblasts highly express CSF1 (Figure S5E) and other growth factors ^51^, which we confirmed by proteomic analysis of the condition media (Figure S5F). Therefore, to test if SASP can promote macrophage proliferation, we treated BMDMs with conditioned media derived from Sen(IR)-fibroblasts and non-senescent fibroblasts and measured cell proliferation with the fluorescent thymidine analogue EdU which is incorporated into DNA during active synthesis (Figure 5J). We observed a 3-fold increase in EdU+ macrophages upon treatment with conditioned medium from Sen(IR)-fibroblasts compared to conditioned medium from non-senescent fibroblasts (Figure 5J). This enhanced proliferation resulted in higher cell confluency after just 24 hours of SASP exposure (Figure 5K).

Finally, since we observed a significant increase in CD38^+^ adipose tissue macrophages in aged eWAT we decided to test whether the SASP from a more adipose tissue specific senescent progenitor cell can also promote CD38 expression in macrophages. Therefore, we focused on pre-adipocytes since they are one of the most abundant progenitor cells that undergo senescence and have recently been implicated in reducing healthspan and lifespan in aging mice^54^. As expected Sen(IR)-preadipocytes, compared to non-irradiated preadipocytes showed elevated B-galactosidase activity (Figure 5L), a hallmark of senescence, and greater expression *p16(Cdkn2a) and p21(Cdkn1a)*, and only modest expression of *Cd38* (Figure 5M). In comparison, BMDMs treated with conditioned medium derived from Sen(IR)-preadipocytes showed significantly elevated *Cd38* expression (Figure 5N). These data provide *in vitro* and *in vivo* evidence that inflammation derived from both microbial and sterile sources of inflammation such as inflammatory cytokines found in the SASP of primary senescent cells is a key driver of CD38 expression in macrophages and influences tissue NAD levels (Figure 5O).

### Aging is associated with increased Cd38 expression by liver Kupffer cells as a result of cellular senescence

Aging is associated with a decline in liver NAD levels, as we observed in Figure 6A, which has recently been reported to be CD38-dependent^14^. However, it is not known what cells in the aged-liver express CD38 and are the key source of CD38 NADase activity. Liver-resident macrophages, known as Kupffer cells, make up to 15% of total cells in the liver and 80-90% of all tissue-resident macrophages ^68^. Activation of Kupffer cells to a pro-inflammatory state has been implicated as a driver of aging-related liver disease such as non-alcoholic liver disease, cirrhosis, fibrosis, but also as a driver of pathogenesis in infectious diseases such as hepatitis^69^. In addition, fat accumulation in the liver in obesity leads to activation of resident Kupffer cells, increased inflammation and liver insulin resistance^68^. Given the connection we found between inflammation, senescence and expression of CD38 by resident adipose tissue macrophages, we investigated whether CD38 expression was significantly altered in Kupffer cells of aged mice livers.

**Figure 6.**
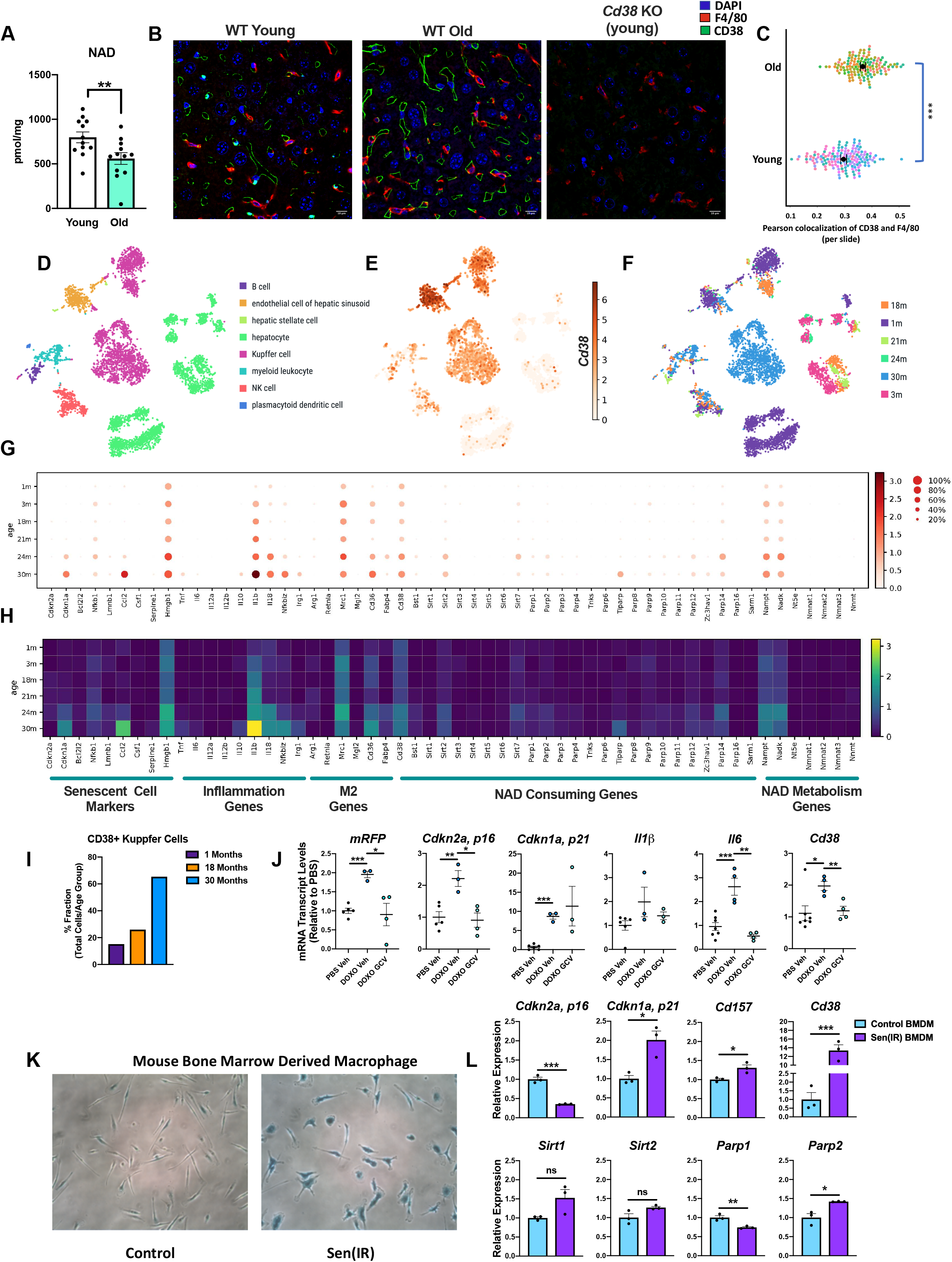
CD38+ Kupffer cells accumulate in the livers of aging mice. (A) LC-MS was used to quantify NAD in the liver from young (4-month) and old (26-month) mice. NAD concentrations are shown as pmol/mg of tissue. (B) Immunofluorescence of the macrophage marker/antigen F4/80 (red), CD38 (Green) and nuclei with DAPI (Blue) in liver from WT young (4-month), WT old (26-month) mice, and young (3-month) *Cd38* KO mouse. Scale bars represents 10μm. Representative of 7-8 mice/group. (C) Analysis of IF images above (a trained neural network to identify macrophage regions and colocalization from Pearson Correlation of F4/80 and CD38) for old and young liver slides (each dot represents one slide), n=20 slides per mouse (7-8 mice/group) (D) Tsne plot of annotated cell populations found in the livers of old and young mice using single-cell transcriptome data from the Tabula-Muris database (https://tabula-muris-senis.ds.czbiohub.org; used for figures D-I). (E) Tsne plot of CD38 expression in liver cell populations in aged mice. (F) Tsne plot of annotated liver cells based on the age of mice. Notice Kupffer cells cluster by age. (G) Dot-plot of the indicated genes in Kupffer cells based on age. (H) Heatmap of the indicated genes in Kupffer cells based on age. (I) Percent fraction of total CD38+ Kupffer cells per total amount of cells per age group. (J) mRNA gene expression of the senescence cell reporter mRFP, senescent cell markers *(Cdkn2a, p16* and *Cdkn1a, p21)*, inflammatory cytokines (*I1β* and *Il6*), and *Cd38* in total liver tissue isolated from p16-3MR mice treated with PBS (vehicle), doxorubicin (DOXO), and DOXO + ganciclovir (GCV). (K) SA-β-Galactosidase staining in control or senescent irradiated (Sen(IR)) BMDMs (day 10). (L) mRNA gene expression of senescence markers (*Cdkn2a, p16* and *Cdkn1a, p21*) and NAD consuming enzymes including *Cd38* in non-senescent control-BMDM or Sen(IR)-BMDMs (day 10). Data is showing the mean ± SEM (n=3 biologically independent experiments). Data from individual mice are shown for *in vivo* experiments. Statistical significance defined as \**P*<0.05, \*\**P*<0.01, and \*\*\**P*<0.00; two-sided Student’s t-test except for 6A one-tailed t-test was used.

Using immunofluorescence to stain for F4/80+ Kupffer cells and CD38 in old and young liver tissues, we observed increased expression of CD38 in liver sinusoids and greater numbers of F4/80+ macrophages near the sinusoids in old vs. young tissue (Figure 6B). Importantly, *Cd38* KO mice showed lack of CD38 staining, validating the specificity of our CD38 antibody (Figure 6B). Since CD38 can also be co-expressed by liver endothelial cells^41^, and since endothelial cells and Kupffer cells are in close proximity to each other in the sinusoid capillary, we used an unbiased trained neural network to analyze our IF images to detect co-expression of CD38 and F4/80. Interestingly, the neural network was able to identify a significant increase in the co-localization of F4/80+ and CD38 (Figure 6C), indicative of greater amounts of CD38+ Kupffer cells in the livers of old mice. To further explore this finding in an unbiased fashion, we collaborated with the Tabula Muris Consortium which recently published a single-cell transcriptomic analysis of aging tissues in mice^70^. Annotation of the single cells of the liver from young and old mice, ranging from 1-30 months revealed eight distinct cell populations including parenchymal cells such as hepatocytes, and non-parenchymal populations such as endothelial, Kupffer cells, and other immune cell populations (Figure 6D). Moreover, our analysis revealed that *Cd38* was primarily expressed in the liver by endothelial cells and Kupffer cells, while other immune cell populations and hepatocytes showed very little *Cd38* expression (Figure 6E). We also noticed that annotated Kupffer cells from aged animals tended to cluster in distinct populations, suggesting significant changes in their pattern of gene expression during aging (Figure 6F). To further analyze *Cd38* gene expression in Kupffer cell, endothelial cells, and hepatocytes populations in aged mice, we measured gene transcript changes in *Cd38*, as well as those of other NAD consuming enzymes, M2 macrophage markers, inflammatory cytokines, and senescent cell markers (Figure 6G and 6H). *Cd38* gene expression was increased in Kupffer cells from old mice (24-30 months), and the number of Kupffer cells that expressed *Cd38* also increased (Figure 6G-I), consistent with our immunofluorescence analysis. Interestingly, increased expression of *Cd38* in aged Kupffer cells was paralleled by increased expression of the nicotinamide salvage pathway enzyme *Nampt*, inflammatory cytokines *Il-1β* and *Il-18*, and the inflammatory transcription factor *Nfkbiz* which is activated downstream of NF-kB, suggesting that Kupffer cells from old mice undergo a pro-inflammatory M1-like polarization. In contrast to Kupffer cells, expression of *Cd38* in endothelial cells of old animals only increased modestly and hepatocytes did not express *Cd38* at any age (Figure S6A and S6B). Thus, similar to our results showing that CD38+ adipose tissue macrophages accumulate in aged eWAT (Figure 4), pro-inflammatory M1-like CD38+ Kupffer cells also increase in the livers of aged mice.

Accumulation of senescent cells in the liver of aged mice has recently been implicated in liver inflammation characterized by increased fat deposition and greater numbers of macrophages in the aged liver^71^. Since increased inflammation and senescence cell burden is sufficient to drive CD38 expression in adipose tissue macrophages, we investigated whether senescent cells also promote CD38 expression in the liver. To test this hypothesis, we used a modified mouse strain that has been extensively utilized in the senescence field. The p16-3MR mouse model is genetically engineered to use the p16 promoter (a gene activated during senescence) to drive the expression of mRFP, renilla luciferase, and HSV-1 thymidine kinase (HSV-TK). This allows both the detection of p16+ senescent cells and a way to selectively kill them by treating the mice with ganciclovir (GSV)^72^. These mice were treated with doxorubicin and, as expected, senescent cell markers *p16(Cdkn2a)* and *p21(Cdkn1a)* increased significantly along with the *p16-mRFP* reporter, indicative of significant accumulation of senescent cells (Figure 6J). Interestingly, this was paralleled by increased CD38 expression, which was reversed after the deletion of senescent cells with GSV (Figure 6J). Thus, the accumulation of senescent cells in the liver of doxo treated mice is both sufficient and necessary to drive enhanced CD38 expression.

Lastly, we unexpectedly noticed a signature of senescence in the transcriptomic data of Kupffer cells in old vs young animals, characterized by increased expression of *p21(Cdkn1a)*, along with enhanced expression of the chemokine *Ccl2,* the DAMP *Hmgb1*, and increased inflammatory cytokine expression (Figure 6G and 6H). While hepatocytes have been reported to be a major source of senescent cells during aging^71^, we did not see a major senescence signature in old hepatocytes (Figure S6A), and it is unknown whether non-parenchymal cells also undergo senescence during aging. Thus, we hypothesized that CD38 expression may occur as a result of the cellular senescence of resident tissue macrophages. To test this hypothesis, we irradiated BMDMs with 10Gy of radiation, which was sufficient to drive both cell cycle arrest and to promote increased SA-Beta-gal activity (Figure 6K), along with a noticeable difference in morphology (Figure 6K). Analysis of gene expression of senescent macrophage revealed a significant increase in *p21(Cdkn1a)* but not *p16(Cdkn2a)*, and a striking increase in the gene expression of *Cd38* (Figure 6L). Thus, senescent macrophages and their resultant SASP may represent a key source of CD38 expression in specific aged tissues such as the liver.

## Discussion

There is growing evidence that NAD levels decline during aging. However, the mechanism is poorly understood. CD38 has recently emerged as a key NAD consuming enzyme that is upregulated during aging. Loss of CD38 activity in *Cd38 KO* mice or in mice treated with a CD38-specific inhibitor, 78c, are protected from age-related NAD decline and show improved metabolic health ^14,41^. Therefore, blocking CD38 NADase activity is emerging as a promising therapeutic target to suppress NAD loss associated with aging ^73^. In this study, we identified macrophages as a key cell type showing increased expression of CD38 during aging and the mechanisms responsible for this enhanced CD38 expression.

We determined that aging-related inflammation drives the polarization of tissue resident macrophages to an M1-like state characterized by the increased expression of CD38 and enhanced consumption of NAD (Figure 1 and 3). As organisms age, DNA damage and genotoxic stress accumulates, leading some cells to undergo cell cycle arrest and senescence. Although senescent cells can play beneficial roles in wound healing and embryogenesis, they also secrete inflammatory factors or SASP that disrupt normal cellular functions, damage tissues, and may promote carcinogenesis ^74,75^. Here, we provide evidence that SASP is also sufficient to increase CD38 expression in macrophages and promote macrophage proliferation, leading to increased accumulation of CD38+ macrophages in both naturally aged mice and in young mice with increased senescent cell burden following treatment with the chemotherapeutic agent doxorubicin (Figure 5 and 6). These results provide a key link between the accumulation of senescent cells in aged tissues and age-related NAD decline. Interestingly, recent studies using senolytics or genetically engineered mouse models (such as p16-3MR and INK-ATTAC mice) have shown that selective killing of senescent cells during aging leads to delayed aging-related diseases and increased healthspan ^52–54,72^. Our data suggest that some of the health benefits derived from eliminating senescent cells may occur as a result of reduced CD38 expression in tissue resident macrophages leading to NAD repletion. Our study also raises new questions on how senescent cells and macrophages communicate and how their interactions influence each other’s function. While we showed that SASP, in particular immunomodulatory cytokines such as IL-6, TNFα, and IL-10, can promote CD38 expression in macrophages (Figure 5H-I), it will be important to thoroughly investigate other components of the SASP to identify specific factors which activate CD38 expression in macrophages. In addition to sterile sources of inflammation such as SASP, we showed that a variety of PAMPs can also promote CD38 expression (Figure 5A). Since aging is associated with a loss of gut epithelial barrier integrity, or “leaky gut” ^49^, endotoxins from commensal bacteria can also synergize with SASP to promote inflammation during aging and/or infection and enhance expression of CD38 (Figure 7). Therefore, future studies will be necessary to better understand and dissociate sterile and non-sterile drivers of inflammation that influence CD38 expression and NAD hydrolysis.

**Figure 7.**
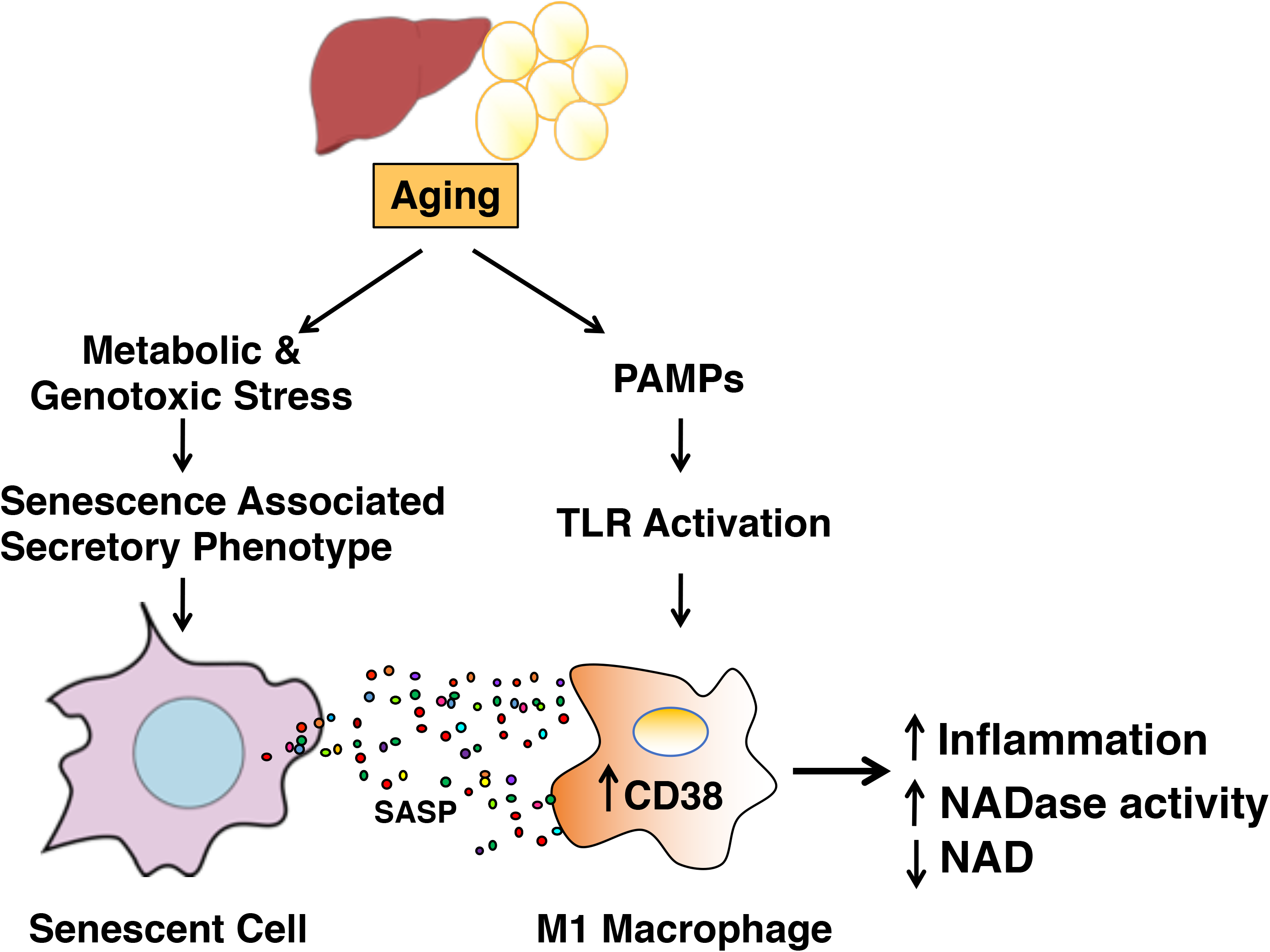
Proposed model how aging-related inflammation leads to enhanced NAD degradation. Aging is associated with the accumulation of DNA damage, which leads to genotoxic stress and the slow accumulation of senescent cells over time. Using *in vivo and in vitro* senescent cell model systems, we show that the accumulation of senescent cells and the accompanying inflammatory cytokines of the senescence-associated secretory phenotype (SASP) is necessary and sufficient to promote CD38 expression and macrophage proliferation. In addition, increased intestinal permeability has been reported to occur during aging leading to enhanced serum levels of endotoxins and other pathogen-associated molecular patterns (PAMPs) which can activate the PRRs expressed by innate immune cells. Collectively, SASP and PAMPs can promote an inflammatory state associated with increased expression of CD38 by tissue-resident M1-like macrophages characterized by enhanced NADase activity.

Previous studies have reported that CD38 regulates immune cell homing, innate immune responses to pathogens, and anti-tumor immunity ^76–78^. However, *Cd38* remains a largely uncharacterized gene in immunology, despite its widespread use in the field as an activation marker. While the teleological purpose of increased NAD consumption by CD38+ M1 macrophages is not obvious, it is tempting to speculate that degradation of NAD may influence NAD levels for control of other NAD consuming enzymes such as PARPs and sirtuins or downstream processes that are dependent on secondary messaging properties of CD38-derived NAD metabolites such as cADPR for control of Ca2+ mobilization ^79^. In addition, NAM exhibits immunomodulatory properties and high levels of NAM can reduce inflammatory cytokine production in macrophages ^80^, suggesting that CD38 may act as a feedback mechanism to initiate the resolution of acute inflammation. This negative feedback mechanism may become maladaptive during aging.

In conclusion, our study demonstrates that increased expression of CD38 in macrophages represents an important source of increased NADase activity during aging. These data also provide insight into how the immune system and NAD metabolism are integrated. While our study focused on metabolic tissues including visceral white adipose tissue and the liver, we speculate that a similar mechanism linking SASP to macrophage CD38 NADase activity may be responsible for NAD declines in other tissues, particularly those where tissue-resident macrophages reside in high abundance such as the brain. This newly discovered cross-talk between senescent cells and macrophages which modulates tissue NAD levels may provide a therapeutic avenue to target age-related inflammation and help restore NAD levels and metabolic health in aging individuals.

## Author Contributions

Conceptualization A.J.C. and E.V.; Methodology, A.J.C. and E.V.; Investigation, A.J.C. (all experiments), A.K. (In vivo experiments and senescent cell experiments), R.P. (NADase assays and flow cytometry), J.A.L.D. (In vivo experiments), A.P. (Single-cell RNA-Seq analysis), H.K. (Flow cytometry), M.S. (LC-MS), C.D.W (In vivo experiments), N. B. (Proteomics), S.I. (RNA analysis), Q.W. (IF imaging and analysis), R. K. (IF imaging and analysis), I.H. (IF imaging and analysis), I.B.S. (LC-MS), M.O. (Animal housing), B.S. (Proteomics), K.I. (Provided *Cd38* and *Cd157* KO mice), S.R.Q. (Single-Cell RNA-Seq Analysis), J.N. (Provided aging mice), C.B. (LC-MS), J.C. (Senescent cell experiments); Writing – Original Draft, A.J.C. and E.V.; Writing – Review & Editing, all authors; Supervision, E.V.; Funding Acquisition, E.V.

## Declaration of Interest

All authors have reviewed and approved the manuscript. C.B. is the inventor of intellectual property on the nutritional and therapeutic uses of NR, serves as chief scientific advisor of and holds stock in ChromaDex. E. V. is a scientific co-founder of NAPA Therapeutics.

## Acknowledgements

This project was supported by the NIH grant R24DK085610 (E.V.), Gladstone Institute intramural funds (E.V.), and Buck Institute intramural funds (E.V.). A.J.C. is a recipient of the UC President’s Postdoctoral Fellowship at UCSF and is also supported by a NIH T32 training grant (3T32AG000266-19S1). A.K. is supported by the SENS Foundation. We thank Marius Walter for his help imaging cells. We thank Vanessa Byles and Davalyn Powell for reviewing the manuscript and helpful discussion.

## Supplementary Materials

**Figure S1**

A) mRNA expression of *CD38* in human peripheral blood monocytes (PBMC) derived macrophages treated with recombinant human IL-4 (M2) or LPS (M1) for 18 hours.

Representative data of one of three patient samples. (n=4 biologically independent experiments)

B) NADase activity measured in human PBMC derived macrophages treated with recombinant human IL-4 (M2) or LPS (M1) for 18 hours. Showing the mean of two separate experiments from different donors with 2 biological replicates for each donor.

C) Schematic of the *de novo* NAD synthesis pathway.

D) mRNA expression of *de novo* NAD synthesis pathway enzymes.

E) Quantification of tryptophan metabolites measured with LC-MS in M0, M2, and M1 mouse BMDMs activated for 24 hours. ND=not detected.

Data is showing the mean ± SEM. n=3 biologically independent experiments except as stated in A and B. Statistical significance defined as \**P*<0.05, \*\**P*<0.01, and \*\*\**P*<0.00; two-sided Student’s t-test.

**Figure S2**

A) mRNA levels of M2 markers in BMDMs pre-treated with or without FK866 and NR for 6 hours prior to stimulation with IL-4 for 16 hours.

B) mRNA levels of M1 markers in BMDMs pre-treated with or without FK866 and NR for 6 hours prior to stimulation with LPS for 6 hours.

Data is showing the mean ± SEM. (n=3 biologically independent experiments). Statistical significance defined as \**P*<0.05, \*\**P*<0.01, and \*\*\**P*<0.00; two-sided Student’s t-test. All statistical comparisons are relative to M2/M1 + FK866.

**Figure S3**

A) Flow cytometry results comparing CD38 surface staining in naive (M0) WT and *Cd38* KO BMDMs or treated with IL-4 (M2) and LPS (M1) for 16 hours.

B) NADase activity measured in intact M0, M2, and M1 WT and *Cd38 KO* BMDMs activated for 16 hours.

C) mRNA expression of *Cd157* in M0 or M1 WT and *Cd38 KO* BMDMs for the indicated times.

D) LC-MS was used to quantify NR in M0 and M1 WT and *Cd38 KO* BMDMs treated for 16 hours.

Data is showing the mean ± SEM. (n=3 biologically independent experiments, n=4 in B).

Statistical significance defined as \**P*<0.05, \*\**P*<0.01, and \*\*\**P*<0.00; two-sided Student’s t-test. Unless noted with a bar, all statistical comparisons are relative to untreated WT or *Cd38* KO sample.

**Figure S4.**

A) mRNA levels of *Il-1α*, *Cxcl1,* and *IL-10* in whole eWAT from 6 month and 25-month-old mice.

B) Flow cytometry gating strategy to identify CD45+ immune cells isolated from the stromal vascular fraction of eWAT. Cells isolated from the SVF of eWAT were first gated on forward scatter (FSCA) vs side scatter (SSCA) to discard cell debris and dead or dying cells. Next FSCH (height) vs FSCA (Area) was used to select for single cells. Single cells were then gated for auto-fluorescent using the Empty (E) BV421 vs BV711 channels (which we did not use as antibody fluorophores) to discard any cells that showed auto-fluorescence in these channels. Next, CD45+ cells were selected and analyzed for CD38 and macrophage markers. Flow cytometry gating strategy to identify resident and non-resident macrophages isolated from the stromal vascular fraction of eWAT, showing representative flow plots and histograms for the indicated age of mice.

C) Flow cytometry quantification of CD38− (low) resident macrophages, CD38-non-resident macrophages, and CD38+ (high) non-macrophage immune cells isolated from eWAT of mice for the indicated age shown.

For *in vivo* experiments, data from individual mice are shown.

Statistical significance defined as \**P*<0.05, \*\**P*<0.01, and \*\*\**P*<0.00; two-sided Student’s t-test.

**Figure S5**

A) mRNA levels of NAD consuming enzymes in whole eWAT from 3-month-old mice IP injected with PBS or LPS for 4-weeks.

B) mRNA levels of *Il-1α*, *Cxcl1,* and *IL-10* in whole eWAT from 6-month-old mice IP injected with doxorubicin or PBS.

C) Quantification of CD38 low resident macrophages, and CD38 low non-resident macrophages isolated from eWAT of 6-month-old mice IP injected with doxorubicin or PBS.

D) CD38 mRNA levels in WT and *Cd38 KO* BMDMs co-cultured (10:1) with non-senescent control mouse dermal fibroblasts (CTRL-MDF) or irradiated senescent MDF (Sen(IR)-MDF) for 24 hours.

E) mRNA levels of inflammatory genes in CTRL-MDF and Sen(IR)-MDF.

F) Heatmap of significantly upregulated proteins identified by mass spectrometry in conditioned media from CTRL-MDF and Sen(IR)-MDF.

For *in vivo* experiments, data from individual mice are shown.

Data is showing the mean ± SEM. (n=4 biologically independent experiments in D). Statistical significance defined as \**P*<0.05, \*\**P*<0.01, and \*\*\**P*<0.00; two-sided Student’s t-test except for 5a and 5b one-tailed t-test was used.

**Figure S6**

A) Dot-plot and heatmap of the indicated genes in liver hepatocytes based on age.

B) Dot-plot and heatmap of the indicated genes in liver endothelial cells based on age.

## Materials and Methods

### Mice

C57BL/6J mice were used for *in vivo* studies and as a source of BMDMs. Mice were maintained at Buck Institute on a standard chow diet and all procedures were performed in accordance with the guidelines set forth by the Institutional Animal Care and Use Committees (IACUC) at the institution. *Cd38* KO mice on a C57BL/6J background were obtained from The Jackson Laboratory (Bar Harbor, ME) and were bred and housed at our animal facility at the Gladstone Institute and Buck Institute. Bone Marrow from mice used in experiments comparing WT, *Cd157* (*Bst1*) KO, *Cd38* KO, and *Cd157/Cd38* DKO were generously provided from the laboratory of Katsuhiko Ishihara.

### BMDM culture and experiments

BMDMs used in experiments were derived from bone marrow extracted from the femurs of euthanized mice (6-12 weeks old male and female C57BL/6J) by mortar and pestle. Briefly, femurs, were placed in the mortar and were washed with 70% ethanol to sterilize followed by two washes with complete RPMI (cRPMI; standard RMPI (Corning) supplemented with 10% fetal calf serum, penicillin-streptomycin solution (Corning), 1mM sodium pyruvate solution (Corning), 2mM L-Glutamine solution (Corning), 10nM HEPES buffer (Corning), and 50μM 2-mercaptoethanol). After washing, 10mls of cRPMI were added to the mortar and the femur bones were gently crushed. The resulting media was collected and filtered through a 70μm filter and placed in a conical tube. The filtered supernatant was centrifuged at 1200 RPM (150 RCF) for 5 minutes. Cells were resuspended, counted and plated at a density of 3×10^6^ cells/10cm dish in 10ml of macrophage growth media (cRPMI containing 25% M-CSF containing L929 conditioned media (made in house)). Cells were left to grow for 7 days to differentiate and were supplemented with 5ml of macrophage growth media on day 5. On day 7, BMDMs (yielding 10-12×10^6^ cells/10cm dish) were lifted off the plate using cold PBS containing 5mM EDTA. BMDMs were counted and replated in macrophage growth media overnight prior to experiments. On day of experiments, macrophage growth media was replaced with cRPMI 6 hours prior to stimulation to remove M-CSF. M2 polarization was performed by stimulating macrophages with 10ng/ml recombinant mouse IL-4 (Peprotech). For M1 polarization macrophages were stimulated with 100ng/ml LPS (LPS EK-Ultrapure, Invivogen). To test how different TLR ligands activate CD38, a Mouse TLR Agonist kit was purchased from Invivogen and BMDMs were treated with each ligand for 16 hours. The NAMPT inhibitor FK866 was purchased from Sigma Aldrich.

### Gene expression measurement

For *in vitro* experiments RNA was isolated using RNA STAT 60 (Amsbio) per manufacturer’s protocol. 1μg RNA was converted to cDNA using a High Capacity cDNA Reverse Transcription Kit (Applied Biosystems), and diluted with 280μl H_2_O post reaction. Gene expression was measured using a CFX384 Real Time System (Bio-Rad) and Thermo Scientific Maxima 2x SYBR Green. Data was analyzed using the CFX Bio-Rad software and normalized using the delta/delta CT method. In vitro BMDMs were normalized to the housekeeping gene hypoxanthine phosphoribosyltransferase (*Hprt*). For in vivo experiments RNA was extracted from flash frozen epididymal fat samples using Trizol (ThermoFisher) in a TissueLyser system (Qiagen). cDNA was generated as described above and gene expression was measured in a LightCycler480 II using the Universal Probe Library system (Roche). Expression was normalized to *Actin* and *Tubulin* using the delta/delta CT method.

### *In vivo* experiments

Male and female C57BL/6J mice were obtained from the National Institute on Aging colony and The Jackson Laboratory (Bar Harbor, ME). Mice ranging in age from 2 to 30-months old were used for determination of the effects of natural aging. 4-to-6-months old mice were injected intraperitoneally with a single dose of 10 mg doxorubicin/kg or vehicle (PBS) and euthanized six weeks later. For elimination of p16-positive cells in the doxorubicin model, p16-3MR mice were i.p. injected with 25 mg/kg ganciclovir (GCV) for 5 consecutive days before euthanasia. For LPS experiments, 8-week old male C57BL/6J mice were obtained from The Jackson Laboratory and were injected with 0.25mg/kg LPS or vehicle (PBS) twice weekly for four weeks (LPS EK-Ultrapure, Invivogen). All animals were euthanized by CO_2_ followed by cervical dislocation. Epididymal fat or liver was excised and portions of the tissue were either fixed for microscopy, flash frozen in liquid nitrogen for qPCR/Western Blot analysis or processed for flow cytometry as described below. All procedures were approved by the Buck Institute IACUC and were in compliance with approved animal protocols.

### NAD Metabolome Mass Spectrometry

For adipose tissue NAD and NADP levels, fat samples were received on dry ice and stored at −80 °C. Samples were pulverized prior to extraction by freezing in LN_2_ and adding to a Bessman pulverizer cooled to LN_2_ temperature. Samples were weighed frozen into microcentrifuge tubes cooled to LN_2_ temperature. The appropriate internal standard solution was added and the sample extracted with hot 75%Ethanol/25% HEPES (pH 7.1) buffer as previously described ^81^. After centrifugation, the aqueous layer between the cell pellet and lipid layer was withdrawn and transferred to a clean microcentrifuge tube; the solvent was evaporated to dryness using a vacuum centrifuge. Standards and controls were prepared by adding internal standard, the appropriate amount of standard working solution, and ethanol/HEPES buffer to microcentrifuge tubes; the standards were dried along with the samples. The dried samples were stored at −20°C until reconstitution immediately prior to the analytical run. For analysis, samples were reconstituted in 70μL of 10mM ammonium acetate and analyzed using 2 separations as previously described^81,82^, and tissue NAD and NADP levels were normalized to tissue weight. Cells Methodology:Briefly, 7×10^6^ BMDMs were cultured in 10cM TC-plates overnight in macrophage growth medium to ∼90% confluence. The following day, the media was replaced with cRPMI 6hrs prior to stimulating with IL-4 (10ng/ml) or LPS (100ng/ml) for the indicated timepoints. BMDMs were washed twice with cold PBS, and gently scraped off, counted, pelleted, flash frozen, and shipped overnight on dry ice to the Brenner Lab. Cell pellets of BMDMs were received on dry ice and stored at −80 °C. To extract metabolites the appropriate internal standard was added along with 400μL of hot ethanol/HEPES buffer as described previously.^1^ Calibrators and controls were prepared and solvent removed in a vacuum centrifuge. The analytical runs were conducted as above. BMDM metabolite levels were normalized to cell number (pmol/1×10^6^ cells) and graphed as normalized peak area. Internal standards:Stable isotope analogs of nucleotides and nucleoside were grown in a yeast broth with universally labelled ^13^C glucose, resulting in all ribose rings being fully labelled. ^13^C_10_ NAD was used as its internal standard. For the second analysis a mix of ^18^O-NR, ^18^O-Nam, d_4_-NA, d_3_-MeNam, and d_3_-methyl-4-pyridone-3-carboxamide was used as internal standard.

### Quantification of tryptophan and metabolites levels

Quantification was performed as previously described ^83^. Briefly, 7×10^6^ BMDMs were cultured in 10cM TC-plates overnight in macrophage growth medium to ∼90% confluence. The following day, the media was replaced with cRPMI 6hrs prior to stimulating with IL-4 (10ng/ml) or LPS (100ng/ml) for 24hrs. Cells were washed twice with cold PBS, and gently scraped off, pelleted, flash frozen, and shipped to the Lee lab on dry ice. The extracted tryptophan and metabolites analyzed by LC-ESI-MS/MS (API 5000 QTRAP mass, AB/SCIEX) by MRM mode. The tryptophan and metabolites MS/MS transitions (*m*/*z*) were 205→188 for tryptophan, 209→192 for kynurenine, 225→208 for 3-hydroxy-L-kynurenine, 154→136 for 3-hydroxyanthranilic acid, 206→160 for kynurenic acid, 190→144 for xanthurenic acid, 138→120 for anthranilic acid, 124→106 for 2-picolinic acid, 168→150 for quinolinic acid, and 199→112 for L-mimosine as an internal standard, respectively. The metabolites were separated on a Luna C18 column (2.1 × 150 mm, 5.0μm) with an injection volume of 5μL and a flow rate of 0.3 ml/min using 0.1% trifluoroacetic acid for mobile phase A and acetonitrile for mobile phase B. The gradient was as follows: 0 min, 2% B; 12 min, 60% B; 13 min, 60% B; 14 min, 2% B; 20 min, 2% B. Data were acquired using Analyst 1.5.1 software (Applied Biosystems, Foster City, CA). Data is presented as pmol/mg protein.

### Extraction of nicotinamide riboside (NR) and analysis by LC-MS/MS

To define the relative abundance of NR by LC-MS/MS analyses, a previously described extraction method optimized for NAD, NADP and NADPH was employed ^84^. Briefly, 7×10^6^ BMDMs were cultured in 10cM TC-plates overnight in macrophage growth medium to ∼90% confluence. The following day, the media was replaced with cRPMI 6hrs prior to stimulating with LPS (100ng/ml) for the indicated timepoints. Cells were washed twice with cold PBS, and gently scraped off, pelleted, flash frozen, and shipped overnight to the Ben-Sahra Lab on dry ice. Metabolites were extracted on ice with 250μl of a 40:40:20 mixture of Acetonitrile/Methanol/ (10mM Tris, pH 9.2, 200mM NaCl). Supernatants were stored at −80°C (one week or less) and 25μl of the cleared solution was injected in a Thermo Q-Exactive (LC-MS) in line with an electrospray source and an Ultimate3000 (Thermo) series HPLC consisting of a binary pump, degasser, and auto-sampler outfitted with a Xbridge Amide column (Waters; dimensions of 4.6mm × 100mm and a 3.5μm particle size). The mobile phase A contained 95% (vol/vol) water, 5% (vol/vol) acetonitrile, 20mM ammonium hydroxide, 20mM ammonium acetate, pH 9.0; B was 100% Acetonitrile. The gradient was as follows: 0 min, 15% A; 2.5 min, 30% A; 7 min, 43% A; 16 min, 62% A; 16-18 min, 75% A; 18-25 min, 15% A with a flow rate of 400μl/min. The capillary of the ESI source was set to 275°C, with sheath gas at 45 arbitrary units, auxiliary gas at 5 arbitrary units and the spray voltage at 4.0 kV. In positive/negative polarity switching mode, an m/z scan range from 70 to 850 was chosen and MS1 data was collected at a resolution of 70,000. The automatic gain control (AGC) target was set at 1×10^6^ and the maximum injection time was 200ms. The top 5 precursor ions were subsequently fragmented, in a data-dependent manner, using the higher energy collisional dissociation (HCD) cell set to 30% normalized collision energy in MS2 at a resolution power of 17,500. The sample volumes of 25μl were injected. Data acquisition and analysis were carried out by Xcalibur 4.0 software and Tracefinder 2.1 software, respectively (both from ThermoFisher Scientific). NR peak area was normalized to protein concentration for each sample and graphed as normalized peak area. Metabolomics services were performed by the Metabolomics Core Facility at Robert H. Lurie Comprehensive Cancer Center of Northwestern University.

### Adipose tissue Digestion

Adipose tissue digestion was performed as previously described ^33^. Briefly, adipose tissue was excised and minced, prior to being digested with 2mg/ml collagenase (Sigma) diluted with KRBH containing 2% FBS for 30 minutes at 37°C. The resulting suspension was filtered through a 70μm filter and centrifuged for 5 minutes at 2000 RPM (425 RCF). The fat layer and supernatant were discarded, and the cell pellet (SVF) was then washed twice with PBS containing 1mM EDTA and 2% FBS prior to lysing red blood cells with ACK lysis buffer (made in house) and staining cells with antibodies for flow cytometry as described below.

### Flow Cytometry

Isolated adipose tissue stromal vascular cells (SVF), cleared of red blood cells using ACK lysis buffer (as described above), were counted and blocked with anti-mouse FC-block (BD Biosciences) per manufacturer’s protocol for 15 minutes on ice. Cells were subsequently labelled on ice for 30 minutes with the following antibodies diluted 1:200; CD38-FITC (clone 90, Biolegend), CD11c-APC (N418, Biolegend), CD11b-PECy7 (M1/70, eBioscience), CD45.2 PE (clone 104, eBioscience), and F4/80-BV510 (clone BM8, Biolegend). Cells were immediately analyzed using the BD FACSAria flow cytometer. Compensation and analysis of the data was performed using the FlowJo software. For in vitro experiments activated BMDMs were lifted off of non-TC treated cell tissue plates after activation using cold PBS with 5mM EDTA for 15 minutes prior to blocking with FC block and staining with antibodies using the protocol described above. Click-iT EdU 488 Flow Cytometry Kit (ThermoFisher) was used to measure macrophage proliferation per manufacturers protocol.

### Immunofluorescent staining and confocal microscopy

Liver and epididymal white fat tissue were dissected and fixed in 4% paraformaldehyde for 24 hours at room temperature followed by paraffin embedding. Paraffin sections (5 μm) were prepared for immunofluorescent staining. After deparaffinization and rehydration, antigen retrieval was performed by boiling slides in 1X Citrate buffer (pH 6.0) using a microwave oven. The slides were then blocked in 10% normal serum from host species of secondary antibody and 5% bovine serum albumin for 1hr in a humidified chamber. Primary antibody incubation was performed at 4°C overnight. The sheep anti-CD38 polyclonal antibody (R&D Systems, Cat# AF4947, diluted at 1:100) and rabbit anti-F4/80 monoclonal antibody (Cell Signaling, Cat# 70076, diluted at 1:100) were used for both single staining and co-staining. The slides were washed 3 times with 1X PBS followed by incubation with fluorescent dye-conjugated secondary antibodies (Jackson ImmunoResearch) at room temperature for 1hr. Following 3 times wash with 1X PBS, slides were mounted with ProLong Gold antifade reagent with DAPI (Thermo Fisher Scientific, Cat# P36935). Images were acquired on a Zeiss LSM700.

### Immunofluorescent Image Analysis

Immunofluorescent images of liver tissue and eWAT were analyzed using a deep convolutional neural network with 23 layers (based on U-Net), trained to detect macrophages in liver tissue and eWAT. For each model, sample images were selected and marked up to identify macrophages (16 samples for Kupffer cells and 12 for macrophages in eWAT). Each training set was extended with data augmentation, where the set of images was automatically and randomly manipulated (by images size, rotation, cropping) to build more robust models. For each model, the neural network was trained for 200 epochs. After training, the neural networks identified 6,019 Kupffer cell regions based on F4/80 staining in 300 slides for 15 mice (8 young at 4 months and 7 old at 26 months). In each region the intensities of CD38 or F4/80 were analyzed, and the Pearson Correlation Coefficient (PCC) was used to measure colocalization.

In addition, a neural network detected 457 macrophage regions in 44 slides of eWAT for 5 young (3 males/2 females) and 4 (2 males/ 2 females) old mice. To compare young and old tissue, the average number of macrophages and the area covered by the macrophages was calculated per slide.

### Human PBMC Macrophage Experiment

Peripheral blood mononuclear cells (PBMCs) were purified from healthy donor blood (Blood Centers of the Pacific, San Francisco, CA) from continuous-flow centrifugation leukophoresis product using density centrifugation on a Ficoll-Paque gradient (GE Healthcare Life Sciences, Chicago, IL). Monocyte were isolated by adherence: 3 ×10^6^ PBMC were seeded into 10cm cell culture dishes and allowed to adhere in a 5% CO_2_ incubator at 37°C for 2-3 hours in cRPMI. Non-adherent cells were removed and the adherent cells were carefully washed, twice with PBS. For the generation of human macrophages, monocytes isolated by adherence were cultured in complete RPMI supplemented with 100ng/ml recombinant human M-CSF (Peprotech) in a 5% CO_2_ at 37°C for 7 days. After 7 days, the cells were stimulated with 100ng/ml LPS or 10ng/ml recombinant human IL-4 for 18 hours.

### ELISA

TNF-α and IL-6 were measured in the media of activated BMDMs using ELISA kits purchased from Biolegend per manufacturer’s protocol. Prior to ELISA supernatants were cleared of debris or dead cells by centrifuging for 5 minutes at 425RCFx5min and diluted 1:150 with the ELISA dilution buffer recommended in the kit.

### Western Blot

Cell lysates were prepared by lysing cells with RIPA Buffer containing Halt protease and phosphatase inhibitor cocktail (ThermoFisher). Protein concentrations were determined using BCA Assay kit (ThermoFisher). Approximately, 15μg of protein lysates were loaded onto 8% polyacrylamide gels and transferred to PVDF membranes. The following primary antibodies were purchased from Cell Signaling and were used to probe for proteins; Tubulin (9F3), Parp1 (46D11), Sirt1 (D1D7), Sirt2 (D4O5O), Sirt3 (D22A3), Sirt5 (D8C3), Sirt6 (D8D12), Sirt7 (D3K5A). CD38 (AF4947) and CD157 (AF4710) antibodies were purchased R&D Systems. Actin (A5441) was purchased from Sigma.

### Arginase Assay

Arginase Assay was performed as previously described ^85^. Briefly, 0.5 ×10^6^ stimulated BMDMs were lysed with 75μl 0.1% TritonX-100 lysis buffer with Halt protease inhibitor cocktail (ThermoFisher). Lysate arginine was activated by adding 50μl of 25mM Tris-HCL and 10μl of 2mM MnCl_2_ to each sample and heating at 56°C for 10 minutes. 100μl of 500mM L-Arginine (pH 9.7) was added to each tube and left to incubate 45 minutes at 37°C. 800μl acid solution (H_2_SO_4_: H_3_PO_4_: H_2_O (1:3:7)) was used to stop each reaction. Urea production was measured by adding 40μl of 9% a-isonitrosopropiophenone (dissolved in 100% ethanol) to each sample and heated to 100°C for 15 minutes. A standard curve of urea was run in parallel to the samples and standards/samples were measured at 540nm using a plate reader.

### NADase Assay

NADase activity was measured using a fluorescence-based assay. Briefly, 0.5 × 10^6^ BMDMs/sample were lysed with 0.1% TritonX-100 Sucrose-Tris (0.25 M sucrose, 40mM Tris [pH 7.4]) lysis buffer with protease and phosphatase inhibitors (ThermoFisher). For the NADase assay on intact macrophages, 2.5 × 10^6^ BMDMs/sample were gently lifted with the cell scraper, centrifuged and resuspended in 100μl PBS/well. The reaction was started adding 80μM of nicotinamide 1,N^6^-etheno-adenine dinucleotide (Sigma Aldrich). The samples were excited at 340 nm and the emission of fluorescence was measured at 460nm at 37°C every min for 1hr in a PHERAstar FS microplate reader (BMG LABTECH). NADase activity was calculated as the slope of the linear portion of the fluorescence-time curve, corrected by the amount of protein in each sample. Protein concentrations were determined using BCA Assay kit (ThermoFisher).

### Senescence Cell and Senescent Cell Condition Media Experiment

Irradiation induced senescence, mouse dermal fibroblasts (MDF) or primary BMDMs were exposed to 10Gy X-rays and medium was changed to fresh medium every 2 days. Control cells were mock irradiated. Primary pre-adipocytes (Pas) were harvested from eWAT of 8-week old C57BL/6J mice as previously described^54^. Briefly, the eWAT was digested as described above, and the resulting SVF was resuspended in complete MEM (Sigma Aldrich) and allowed to adhere to tissue culture plates overnight. The next day, attached cells were washed with PBS and trypsinized. The trypsinized cells, ∼85% PAs confirmed by flow cytometry, were replated and serial passaged. PAs were exposed to 10Gy X-rays and medium was changed to fresh medium every 2 days. Control cells were mock irradiated and serial passaged in parallel up to Day 10 post radiation at which point both irradiated and mock PA conditioned media and cells were harvested. For chemotherapy-induced senescence MDF were treated with doxorubicin (Sigma Aldrich) at concentration 250nM in DMSO for 24 hr. The medium was replaced by normal DMEM supplemented with 10% FBS and refreshed every 2 days. Control cells were treated with equal amount of DMSO. After day 10 of treatment, conditioned medium (CM) were prepared by washing cells 3 times in PBS, then incubating in media for 24h. Bone marrow derived macrophages then incubated with this CM for 24 hours and harvested for RNA isolation. For co-culture experiments, 1×10^6^ WT or *Cd38 KO* BMDMs were counted and added to 6-well tissue culture plates containing 1×10^4^ IRR MDFs or control MDFs at a ratio of (10:1) in 3.5mls of DMEM supplemented with 10% FBS in a 6-well plate for 24 hours. Co-culture cells were harvested with RNA Stat 60 to analyze gene expression of bulk RNA from both macrophages and senescent cells as described above.

### Proteomic Sample Preparation

#### Chemicals

Acetonitrile (#AH015) and water (#AH365) were obtained from Burdick #x0026; Jackson (Muskegon, MI). Reagents for protein chemistry including iodoacetamide (IAA, #I1149), dithiothreitol (DTT, #D9779), formic acid (FA, #94318-50ML-F), and Triethylammonium bicarbonate buffer 1.0 M, pH 8.5±0.1 (#T7408) were purchased from Sigma Aldrich (St. Louis, MO). Urea (#29700) was purchased Thermo Scientific (Waltham, MA). Sequencing grade trypsin (#V5113) was purchased from Promega (Madison, WI). HLB Oasis SPE cartridges (#186003908) were purchased from Waters (Milford, MA).

Induction of Senescence

#### X-ray irradiation

Induction of senescence by ionizing radiation (IR) was performed on mouse dermal fibroblasts with 10 Gy X-ray. Quiescent control cells were mock irradiated. Senescent cells were cultured for 10 days to allow development of the senescent phenotype, and quiescent cells were cultured in 0.2% serum for 3 days. Subsequently, cells were washed with PBS (Gibco #10010-023) and placed in serum-free and phenol-red-free DMEM (Gibco #21063-029) and conditioned media was collected after 24 hours to obtain the secreted protein fractions.

#### Isolation, concentration, and quantification of secreted proteins

Proteins secreted into serum-free medium over a 24-hr period were collected from senescent and quiescent (control) mouse dermal fibroblasts. Samples were concentrated using Amicon Ultra-15 Centrifugal Filter Units with 3 kDa molecular weight cutoff (MilliporeSigma #UFC900324) per manufacturer instructions and transferred into a solution of 8M urea/50 mM Triethylammonium bicarbonate buffer at pH 8. Protein quantitation was then performed using a BCA Protein Assay Kit (Pierce #23225, Waltham, MA).

#### Digestion

An aliquot of 25-100 μg from each sample was then brought to equal volume with 50 mM Triethylammonium bicarbonate buffer in 8M urea at pH 8. The protein mixtures were reduced with 20 mM DTT (37°C for 1 hour), and subsequently alkylated with 40 mM iodoacetamide (30 minutes at RT in the dark). Samples were diluted 10-fold with 50 mM Triethylammonium bicarbonate buffer at pH 8 and incubated overnight at 37°C with sequencing grade trypsin (Promega) added at a 1:50 enzyme:substrate ratio (wt/wt).

#### Desalting

The peptide supernatants were then collected and desalted with Oasis HLB 30 mg Sorbent Cartridges (Waters #186003908, Milford, MA), concentrated, and re-suspended in a solution containing mass spectrometric ‘Hyper Reaction Monitoring’ peptide standards (HRM, Biognosys #Kit-3003, Switzerland) and 0.2% formic acid in water.

##### Mass Spectrometry Analysis

Samples were analyzed by reverse-phase HPLC-ESI-MS/MS using the Eksigent Ultra Plus nano-LC 2D HPLC system (Dublin, CA) combined with a cHiPLC System, which was directly connected to a quadrupole time-of-flight SCIEX TripleTOF 6600 or a TripleTOF 5600 mass spectrometer (SCIEX, Redwood City, CA). Typically, mass resolution in precursor scans was ∼ 45,000 (TripleTOF 6600), while fragment ion resolution was ∼15,000 in ‘high sensitivity’ product ion scan mode. After injection, peptide mixtures were transferred onto a C18 pre-column chip (200 μm x 6 mm ChromXP C18-CL chip, 3 μm, 300 Å, SCIEX) and washed at 2 μl/min for 10 min with the loading solvent (H_2_O/0.1% formic acid) for desalting. Subsequently, peptides were transferred to the 75 μm x 15 cm ChromXP C18-CL chip, 3 μm, 300 Å, (SCIEX), and eluted at a flow rate of 300 nL/min with a 3 h gradient using aqueous and acetonitrile solvent buffers.

For label-free relative quantification all study samples were analyzed by data-independent acquisitions (DIA), or specifically variable window SWATH acquisitions. In these SWATH acquisitions, instead of the Q1 quadrupole transmitting a narrow mass range through to the collision cell, windows of variable width (5 to 90 m/z) are passed in incremental steps over the full mass range (m/z 400-1250). The cycle time of 3.2 sec includes a 250 msec precursor ion scan followed by 45 msec accumulation time for each of the 64 SWATH segments. The variable windows were determined according to the complexity of the typical MS1 ion current observed within a certain m/z range using a SCIEX ‘variable window calculator’ algorithm (i.e. more narrow windows were chosen in ‘busy’ m/z ranges, wide windows in m/z ranges with few eluting precursor ions) ^1^. SWATH MS2 produces complex MS/MS spectra which are a composite of all the analytes within each selected Q1 m/z window. This method requires high scan speeds and high-resolution capabilities, as well as a ‘reference spectral library’ that can be either generated in-house or taken from existing comprehensive and large-scale studies. All collected mass spectrometry data was processed in Spectronaut using an in-house spectra library generated by data-dependent analysis. Data-dependent acquisitions (DDA) were carried out on eight samples containing secreted proteins from mouse dermal fibroblasts to obtain MS/MS spectra for the 30 most abundant precursor ions (100 msec per MS/MS) following each survey MS1 scan (250 msec), yielding a total cycle time of 3.3 sec. For collision induced dissociation tandem mass spectrometry (CID-MS/MS), the mass window for precursor ion selection of the quadrupole mass analyzer was set to ± 1 m/z using the Analyst 1.7 (build 96) software.

##### Quantification and statistical analysis

To build a spectral library, raw files obtained by DDA analysis of eight secreted protein samples were searched using ProteinPilot software (SCIEX), and search results were then imported into Spectronaut software (Biognosys) to create a spectral library. SWATH acquisitions were quantitatively processed using the proprietary Spectronaut v12 (12.020491.3.1543) software^2^ from Biognosys. Quantitative SWATH MS2 data analysis was based on extracted ion chromatograms (XICs) of 6-10 of the most abundant fragment ions in the identified spectra.

Relative quantification was performed comparing different conditions (senescent versus control) assessing fold changes for proteins from the investigated cells and conditions. The number of biological replicates analyzed by proteomics were five senescent secretome samples and six quiescent (control) secretome samples. Significance was assessed using FDR corrected q-values<0.05. See Data Availability (below) for quantitative results from mass spectrometric SWATH analysis.

##### Data availability

All raw files are uploaded to the Center for Computational Mass Spectrometry, MassIVE, and can be downloaded using the following ftp link ftp://massive.ucsd.edu/MSV000083726 (MassIVE ID number: MSV000083726). Data uploads include the protein identification and quantification details, spectral library, and FASTA file used for mass spectrometric analysis.

MassIVE: MSV000083726 (ftp://massive.ucsd.edu/MSV000083726)

## Statistical Analysis

Statistical analysis was performed using GraphPad Prism software and Microsoft Excel. One-sided and two-sided Student’s t-tests were applied to comparisons between two conditions with mean ± SEM shown. NS, not significant, \**P*<0.05, \*\**P*<0.01, and \*\*\**P*<0.001. For statistical analysis of proteomic data please see the methods section above.

