## Supplementary Figures for "Aging-related inflammation driven by cellular senescence enhances NAD consumption via activation of CD38^+^ pro-inflammatory macrophages"

### Sup Figure 1

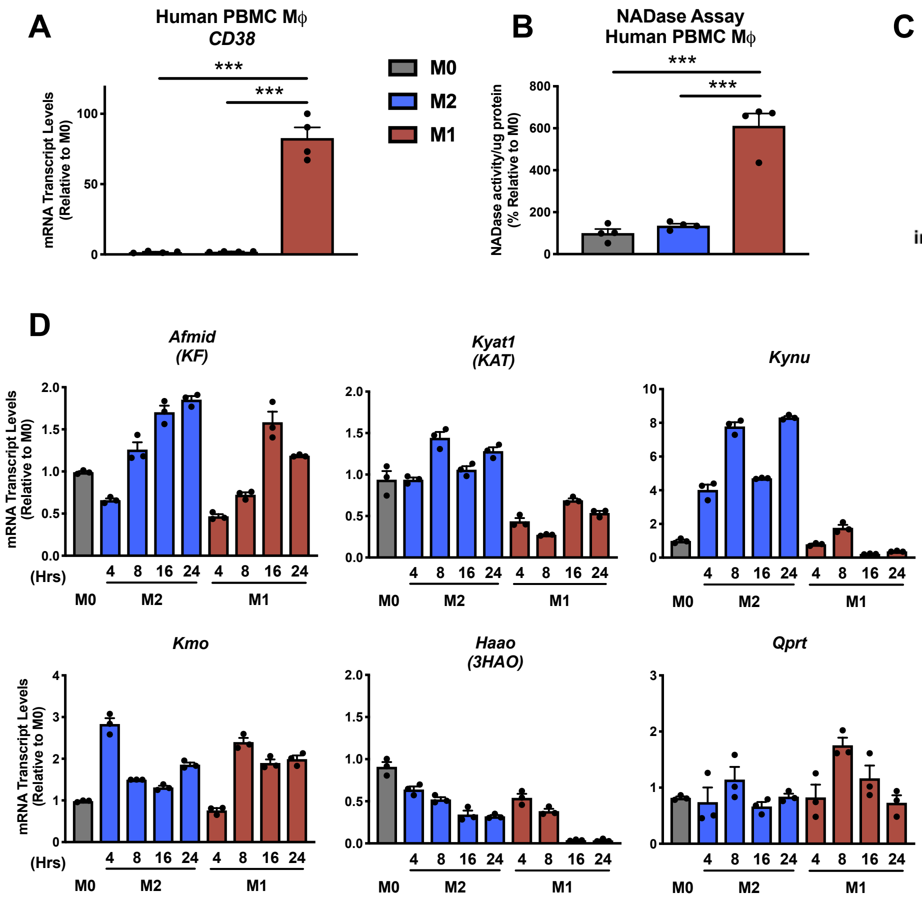


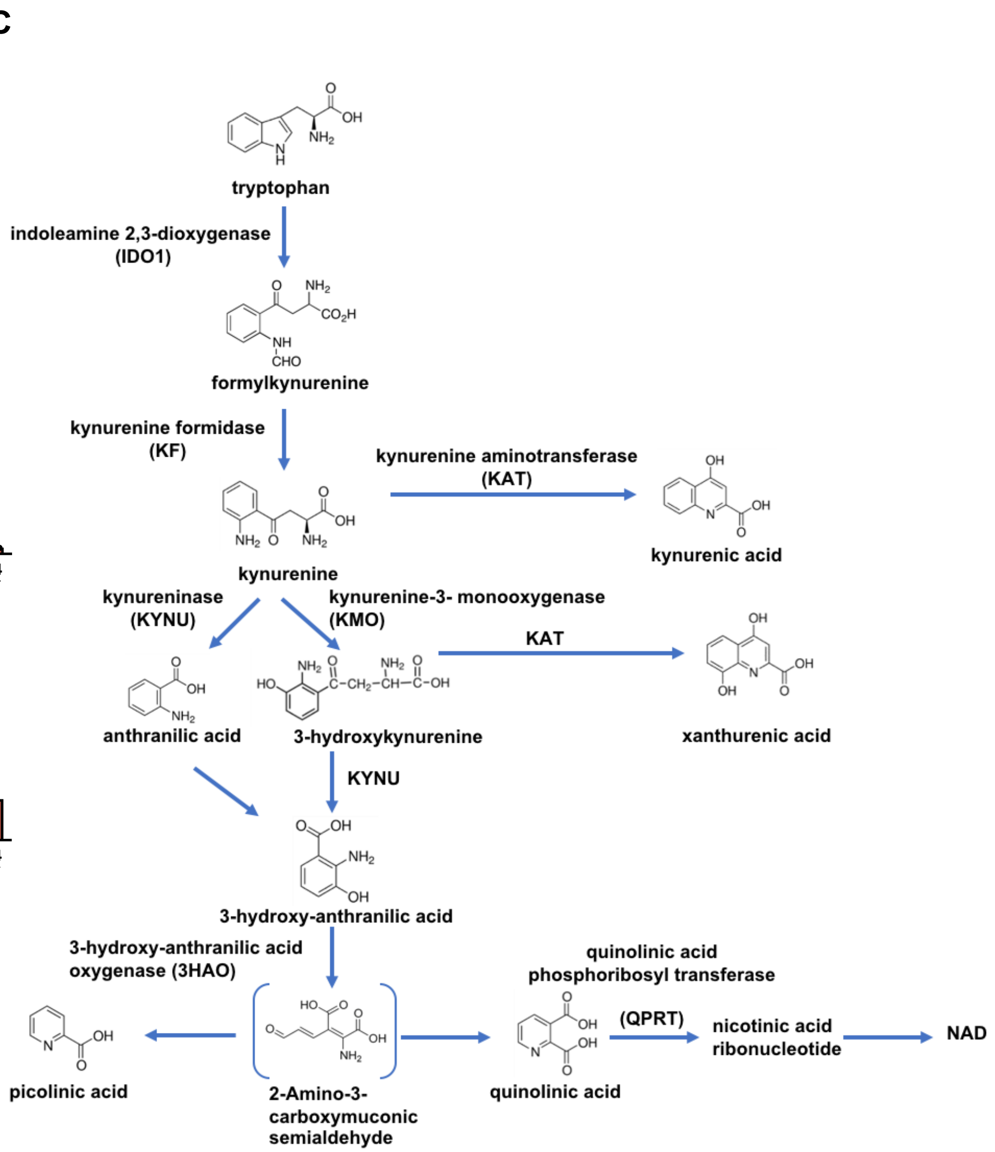


**E** unit: pmol/mg protein

| Macrophage Sample (24 hrs) | Tryptophan | Kynurenine | 3-hydroxy-L-kynurenine | 3-Hydroxy-anthranilic acid | Kynurenic acid | Xanthurenic acid | Anthranilic acid | Picolinic acid | Quinolinic acid |
|---|---|---|---|---|---|---|---|---|---|
| M0 | 25.73 | 0.28 | ND | ND | 2.21 | ND | ND | ND | ND |
| M0 | 24.69 | 0.26 | ND | ND | 2.11 | ND | ND | ND | ND |
| M0 | 24.57 | 0.25 | ND | ND | 2.3 | ND | ND | ND | ND |
| M2 | 9.43 | 0.17 | ND | ND | 0.84 | ND | ND | ND | ND |
| M2 | 11.08 | 0.19 | ND | ND | 0.96 | ND | ND | ND | ND |
| M2 | 9.37 | 0.39 | ND | ND | 0.86 | ND | ND | ND | ND |
| M1 | 640 | 10.71 | ND | ND | 42.89 | ND | ND | ND | ND |
| M1 | 523.94 | 9.42 | ND | ND | 43.94 | ND | ND | ND | ND |
| M1 | 603.24 | 7.01 | ND | ND | 51.57 | ND | ND | ND | ND |

### Sup Figure 2

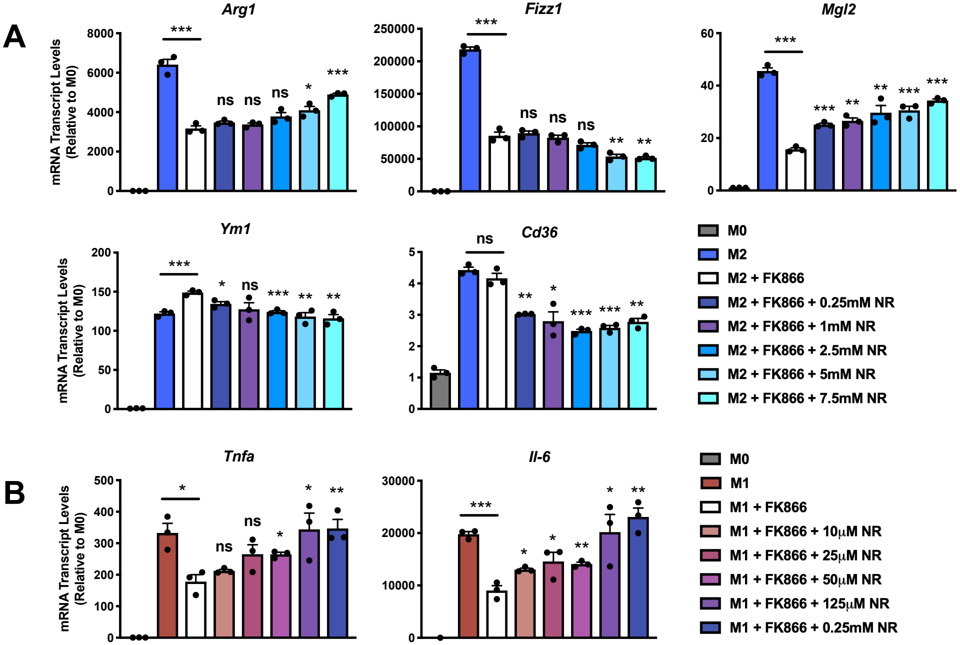
A
Arg1
Fizz1
Mgl2
Ym1
Cd36
mRNA Transcript Levels (Relative to M0)
ns
***
**
*
M0
M2
M2 + FK866
M2 + FK866 + 0.25mM NR
M2 + FK866 + 1mM NR
M2 + FK866 + 2.5mM NR
M2 + FK866 + 5mM NR
M2 + FK866 + 7.5mM NR
B
Tnfa
Il-6
M0
M1
M1 + FK866
M1 + FK866 + 10μM NR
M1 + FK866 + 25μM NR
M1 + FK866 + 50μM NR
M1 + FK866 + 125μM NR
M1 + FK866 + 0.25mM NR

**Sup Figure 3**

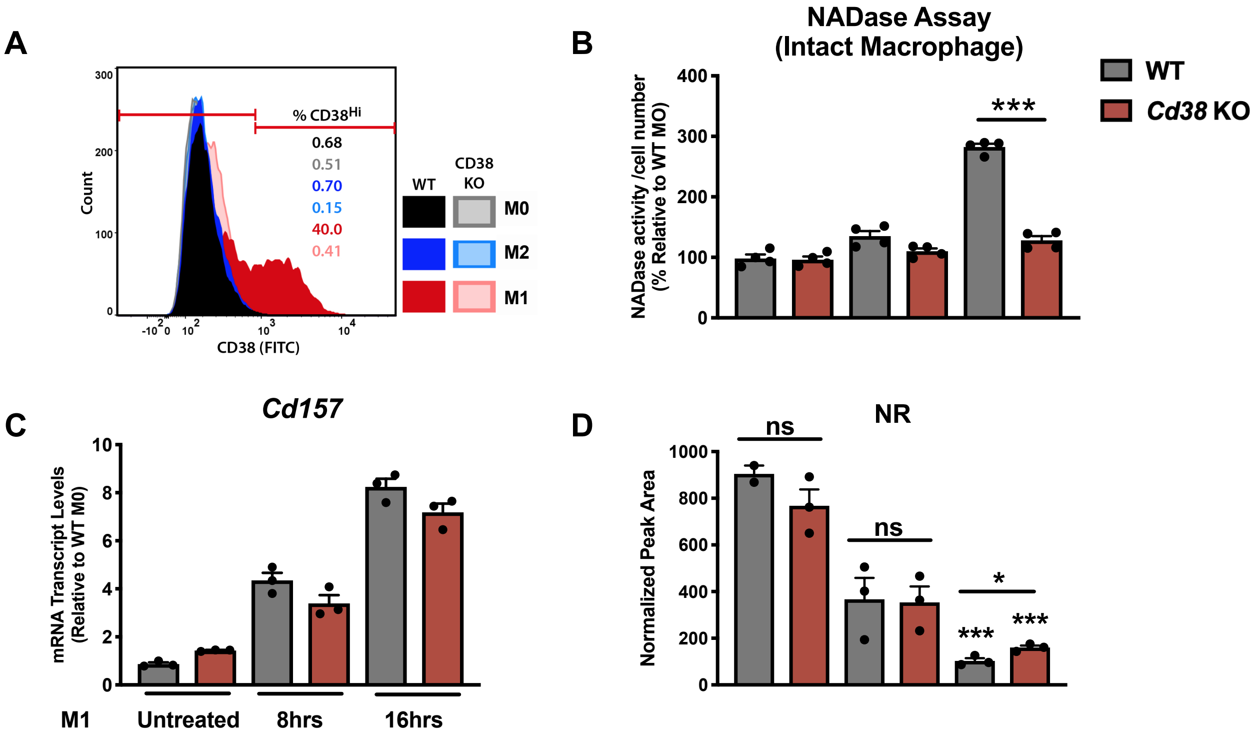

A
300
200
100
0
Count
% CD38Hi
0.68
0.51
0.70
0.15
40.0
0.41
-10^2 0 10^2 10^3 10^4
CD38 (FITC)
WT
CD38 KO
M0
M2
M1
B
NADase Assay
(Intact Macrophage)
WT
Cd38 KO
400
300
200
100
0
NADase activity /cell number
(% Relative to WT MO)
***
C
Cd157
10
8
6
4
2
0
mRNA Transcript Levels
(Relative to WT M0)
M1
Untreated
8hrs
16hrs
D
NR
1000
800
600
400
200
0
Normalized Peak Area
ns
ns
*
***
***

#### Sup Figure 4

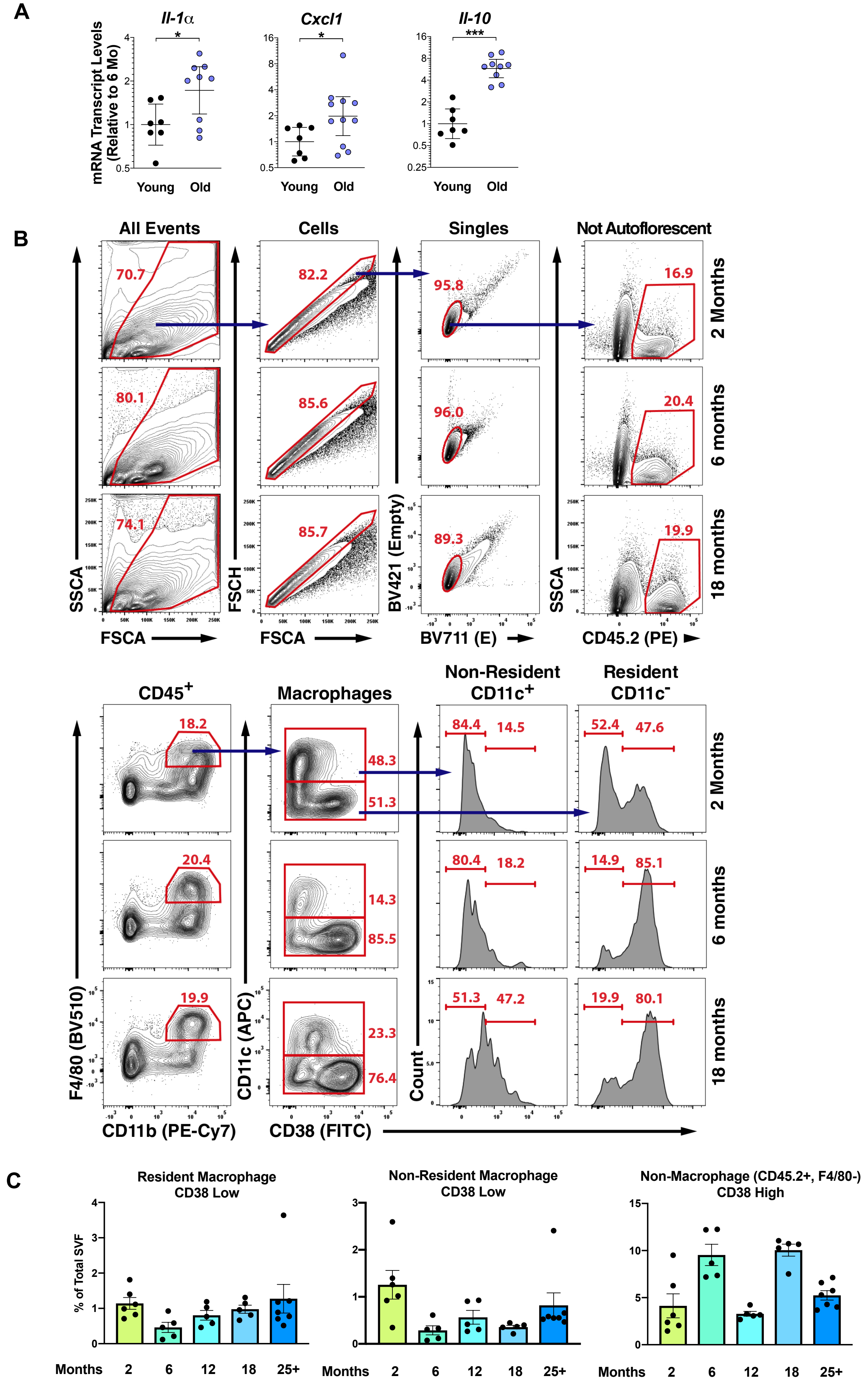

A
Il-1α
Cxcl1
Il-10
mRNA Transcript Levels (Relative to 6 Mo)
*
***
Young
Old
B
All Events
Cells
Singles
Not Autoflorescent
70.7
82.2
95.8
16.9
80.1
85.6
96.0
20.4
74.1
85.7
89.3
19.9
2 Months
6 months
18 months
SSCA
FSCA
FSCH
BV421 (Empty)
BV711 (E)
CD45.2 (PE)
CD45+
Macrophages
Non-Resident CD11c+
Resident CD11c-
18.2
48.3
51.3
84.4
14.5
52.4
47.6
20.4
14.3
85.5
80.4
18.2
14.9
85.1
19.9
23.3
76.4
51.3
47.2
19.9
80.1
F4/80 (BV510)
CD11b (PE-Cy7)
CD11c (APC)
CD38 (FITC)
Count
C
Resident Macrophage CD38 Low
Non-Resident Macrophage CD38 Low
Non-Macrophage (CD45.2+, F4/80-) CD38 High
% of Total SVF
Months
2
6
12
18
25+

### Sup Figure 5

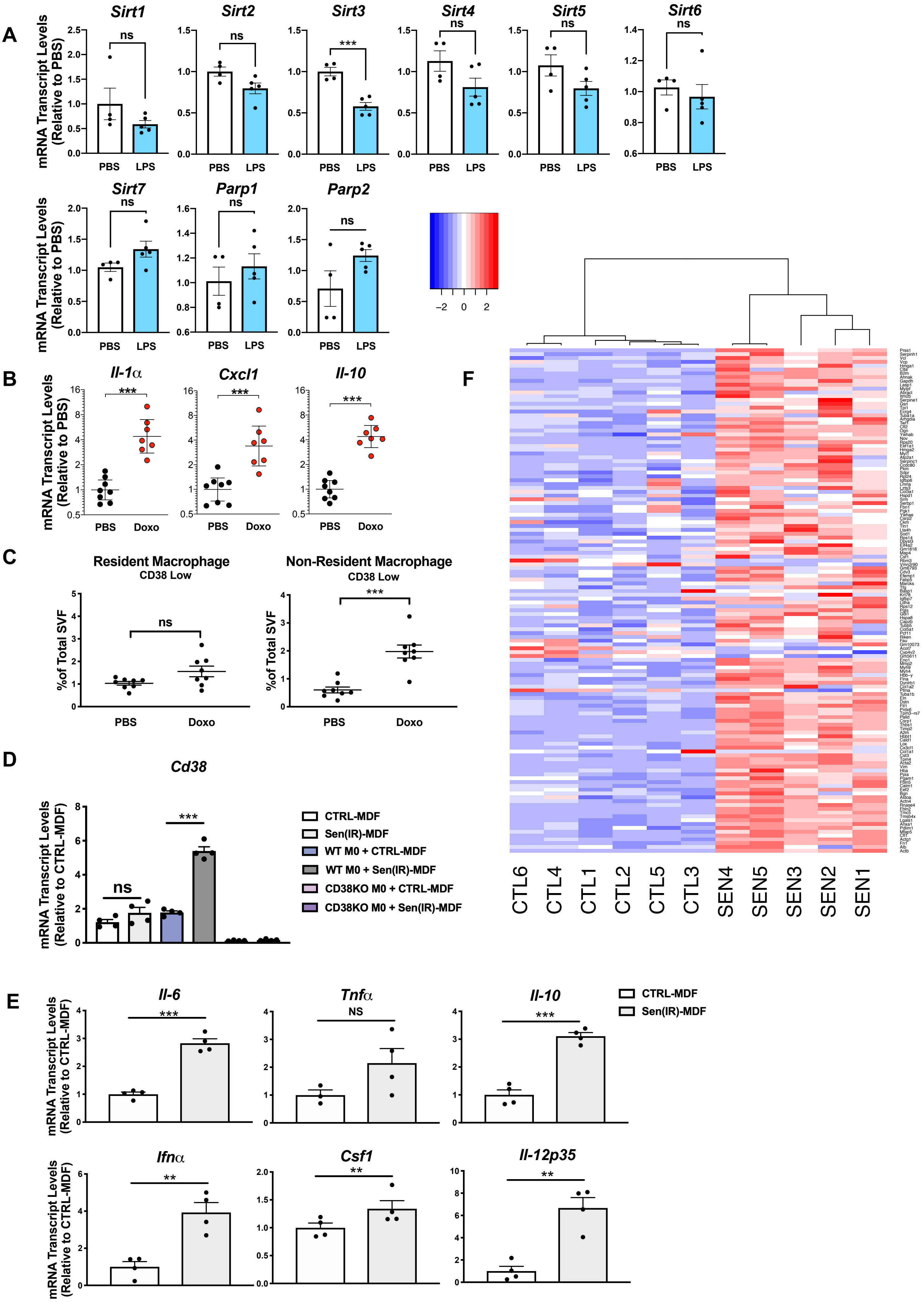

A
Sirt1
Sirt2
Sirt3
Sirt4
Sirt5
Sirt6
Sirt7
Parp1
Parp2
ns
***
mRNA Transcript Levels (Relative to PBS)
PBS
LPS
−2
0
2
B
Il-1α
Cxcl1
Il-10
Doxo
C
Resident Macrophage
CD38 Low
Non-Resident Macrophage
CD38 Low
%of Total SVF
D
Cd38
mRNA Transcript Levels (Relative to CTRL-MDF)
CTRL-MDF
Sen(IR)-MDF
WT M0 + CTRL-MDF
WT M0 + Sen(IR)-MDF
CD38KO M0 + CTRL-MDF
CD38KO M0 + Sen(IR)-MDF
E
Il-6
Tnfα
NS
Il-10
Ifnα
Csf1
Il-12p35
**
F
CTL6
CTL4
CTL1
CTL2
CTL5
CTL3
SEN4
SEN5
SEN3
SEN2
SEN1

### Sup Figure 6

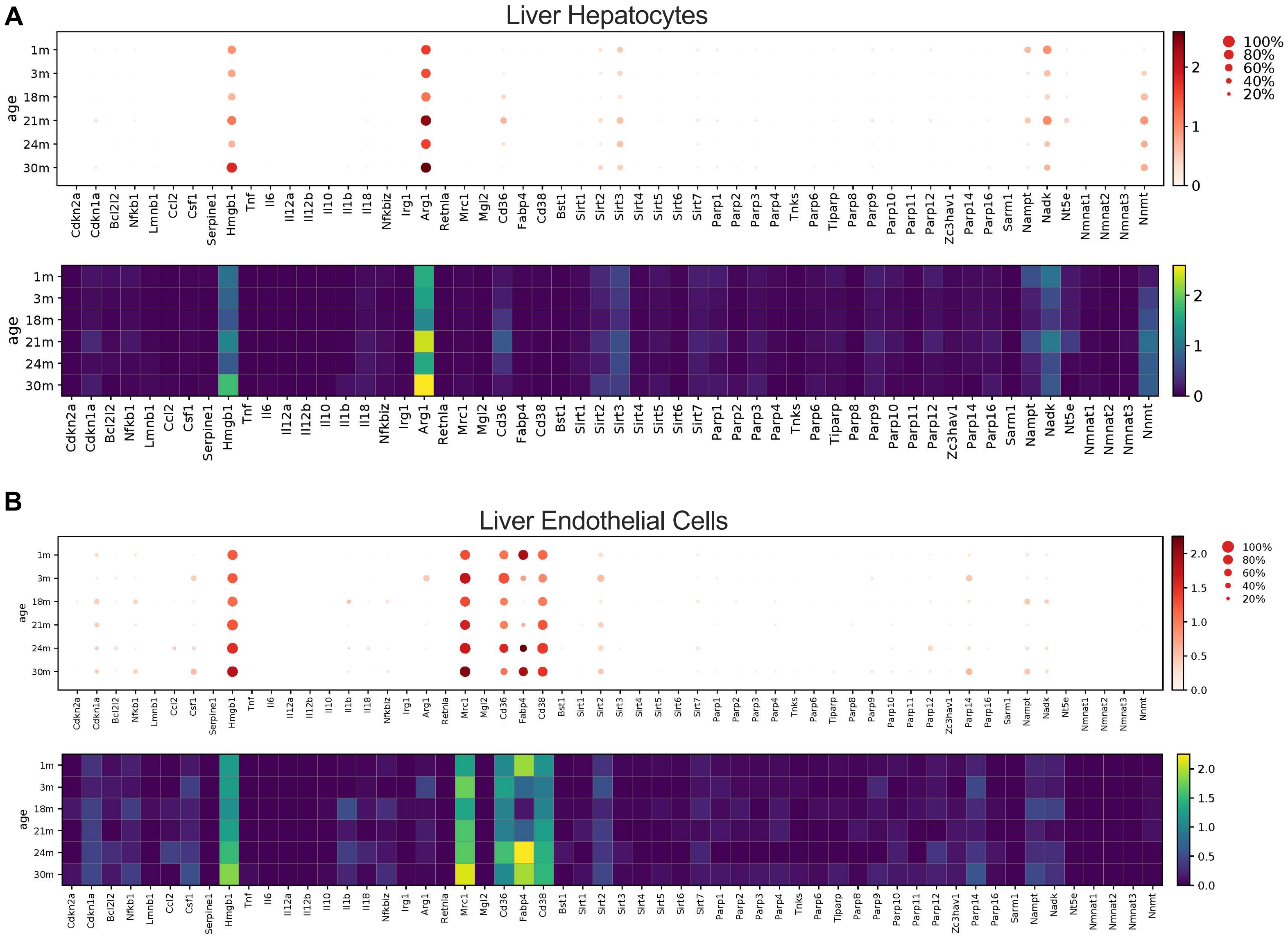

A
Liver Hepatocytes
B
Liver Endothelial Cells
age
1m
3m
18m
21m
24m
30m
100%
80%
60%
40%
20%
0
1
2
0.0
0.5
1.0
1.5
2.0
Cdkn2a
Cdkn1a
Bcl2l2
Nfkb1
Lmnb1
Ccl2
Csf1
Serpine1
Hmgb1
Tnf
Il6
Il12a
Il12b
Il10
Il1b
Il18
Nfkbiz
Irg1
Arg1
Retnla
Mrc1
Mgl2
Cd36
Fabp4
Cd38
Bst1
Sirt1
Sirt2
Sirt3
Sirt4
Sirt5
Sirt6
Sirt7
Parp1
Parp2
Parp3
Parp4
Tnks
Parp6
Tiparp
Parp8
Parp9
Parp10
Parp11
Parp12
Zc3hav1
Parp14
Parp16
Sarm1
Nampt
Nadk
Nt5e
Nmnat1
Nmnat2
Nmnat3
Nnmt
